# Tissue context determines the penetrance of regulatory DNA variation

**DOI:** 10.1101/2020.06.27.175422

**Authors:** Jessica M. Halow, Rachel Byron, Megan S. Hogan, Raquel Ordoñez, Mark Groudine, M. A. Bender, John A. Stamatoyannopoulos, Matthew T. Maurano

## Abstract

Assessment of the functional consequences of disease-associated sequence variation at non-coding regulatory elements is complicated by their high degree of context sensitivity to both the local chromatin and nuclear environments. Allelic profiling of DNA accessibility across individuals has shown that only a select minority of sequence variation affects transcription factor (TF) occupancy, yet the low sequence diversity in human populations means that no experimental assessment is available for the majority of disease-associated variants. Here we describe high-resolution *in vivo* maps of allelic DNA accessibility in liver, kidney, lung and B cells from 5 increasingly diverged strains of F1 hybrid mice. The high density of heterozygous sites in these hybrids enables precise quantification of the effect size and cell-type specificity of hundreds of thousands of variants throughout the mouse genome. We show that chromatin-altering variants delineate characteristic sensitivity profiles for hundreds of TF motifs. We develop a compendium of TF-specific sensitivity profiles accounting for genomic context effects. Finally, we link these maps of allelic accessibility to allelic transcript levels in the same samples. This work provides a foundation for quantitative prediction of cell-type specific effects of non-coding variation on TF activity, which will dramatically facilitate both fine-mapping and systems-level analyses of common disease-associated variation in human genomes.

## Introduction

Systematic census of cis-regulatory elements using genome-wide profiling of DNA accessibility to the endonuclease deoxyribonuclease I (DNase I) has critically informed understanding of tissue-specific gene regulation^1^and the genetics of common human diseases and traits^2^. But these maps provide only indirect evidence for the function of regulatory DNA and cannot address the effects of sequence variation therein. Regulatory element function depends on both genomic and cellular context, which cannot be easily recapitulated in reporter assays^3^. Profiling of DNA accessibility or protein occupancy at polymorphic sites represents a genome-scale approach to assessing local effects of regulatory variation in context^4-8^. However, this approach is limited by low sequence diversity in an individual human genome and the difficulty of accessing many disease-relevant cell types. Recognition of functional human sequence variants has thus been impeded by the lack of large-scale datasets assaying function at their endogenous context in vivo.

The laboratory mouse Mus musculus and related species have long been a key model for human disease and genome function^9,10^. Given the near-complete conservation of transcriptional regulatory machinery with humans, mouse transgenic experiments have been foundational in the understanding of human genetics and gene regulation^11,12^. The availability of mice from divergent strains/species offers a rich trove of genetic diversity dramatically exceeding that in human populations^9^, and with potential access to a variety of tissues and cell types including developmental timepoints^13^. Genomic approaches have linked many of these DNA sequence changes to altered transcription factor (TF) binding^14,15^, chromatin features^16,17^, gene expression^18-20^, and protein levels^21^, and further dissection of molecular traits is highly complementary to high-throughput knockout phenotyping studies^22,23^.

DNase I-hypersensitive site (DHS) maps in mouse tissues show substantial divergence in regulatory DNA compared to human DHSs^2,24^, suggesting that studies of human cis-regulatory variation cannot directly incorporate analyses of orthologous mouse loci. Past work has shown that genetic effects on chromatin features can be modeled using TF-centric analysis^4,5^. The high conservation of trans-regulatory circuitry suggests that such a TF-centric approach might be able to leverage the power of mouse genetics for interpretation of human cis-regulatory variation.

Here we describe high-resolution maps of allelic DNA accessibility in 4 cell and tissue types across a series of F1 hybrid mice derived from inbred lab- and wild-derived strains and species. These maps reveal genetic effects on DNA accessibility which are moderated by cell and tissue context. We use these maps to derive sensitivity profiles for hundreds of TFs, facilitating prediction of functional noncoding polymorphism across mammalian genomes. Finally, we use matching RNA-seq data to assess the correlation between accessibility and expression levels.

## Results

### Allelic analysis of DNA accessibility

We analyzed hybrid, fully heterozygous F1 mice resulting from a cross of the reference C57BL/6J with five diverged strains or species: 129S1/SvImJ, C3H/HeJ, CAST/EiJ, PWK/PhJ, and SPRET/EiJ. We mapped DHSs in four diverse cell and tissue types, including whole kidney, liver, lung, and B cells purified from femoral bone marrow (**Fig. 1a-b**). We selected the highest-quality samples for deep paired-end Illumina sequencing based on fragment length distribution (**Fig. 1c, Supplementary Fig. 1**) and high signal-to-noise demonstrated by a mean Signal Portion of Tags (SPOT) score of 60% (**Supplementary Table 1**). A total of 67 samples were sequenced to an average of 203M reads each, including at least 2 replicates per condition (median = 3 replicates) (**Supplementary Table 1**). We developed a stringent mapping procedure requiring high mappability to both the reference and a customized strain-specific genome incorporating known single nucleotide variants (SNVs) and indels^22^ (Methods). Replicate samples exhibited a median correlation in DNaseI cleavage density at DHSs of 0.93 (**Supplementary Fig. 2**).

**Fig. 1.**
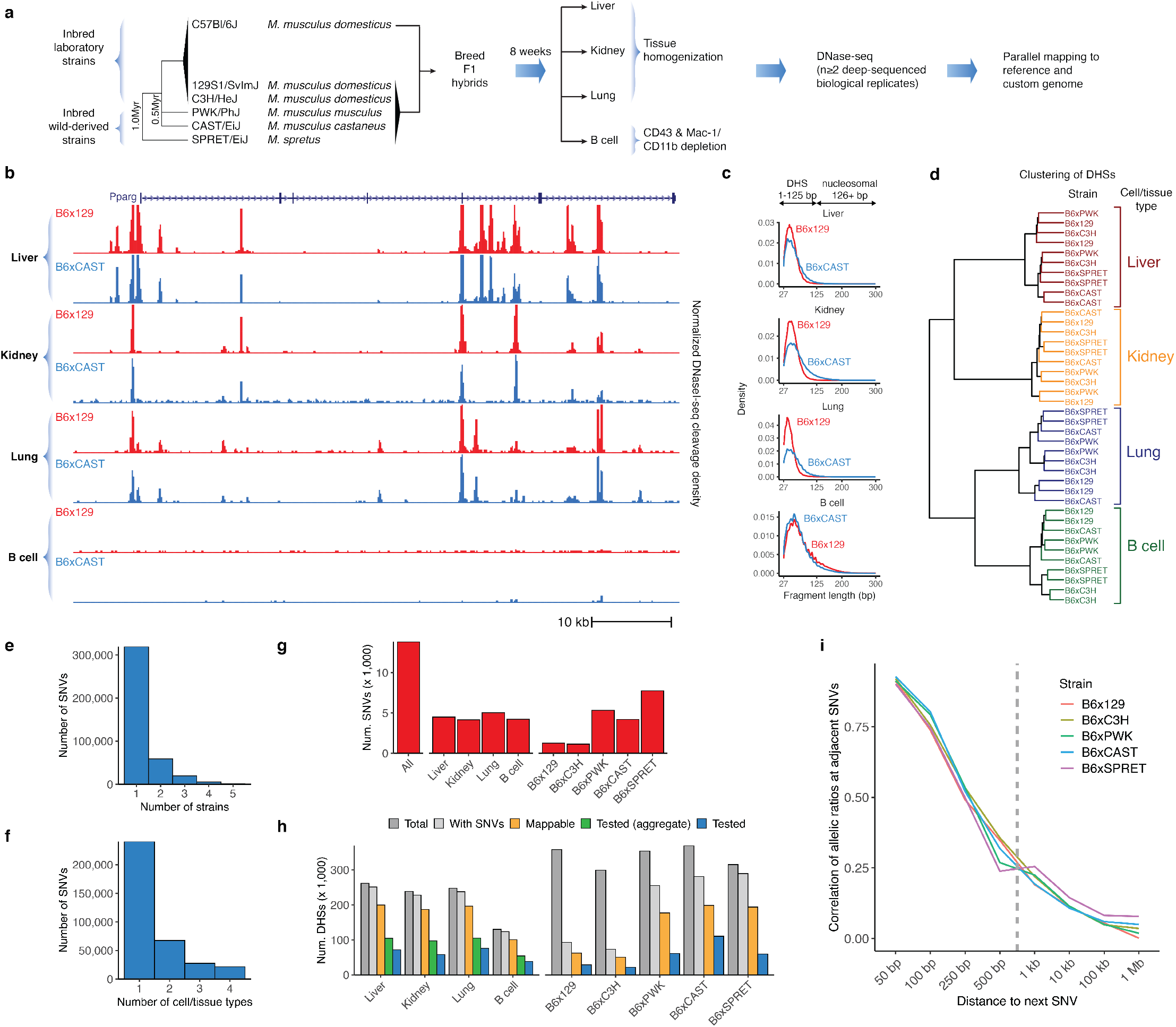
Allelic analysis of DNA accessibility in hybrid mice from diverged strains. **a**. Overall schematic of experiment **b**. DNase-seq profiles at the Pparg locus in liver, kidney, lung tissue and B cells from F1 crosses of C57Bl/6J dams with 129/SvImJ and CAST/EiJ sires. **c**. Fragment length distribution of samples showing high-quality libraries comprising non-nucleosomal fragments. **d**. Hierarchical clustering of DHSs from high-depth samples. **e**.-**f**. Counts of SNVs shared across strains (**e**) and cell types (**f**). **g**. Counts of imbalanced SNVs (FDR 10%). Counts are reported in aggregate across all data sets (left), by cell type (middle), and by parental strain (right). **h**. Summary of master list of DHSs overlapping SNVs from all strains. Counts include all DHSs (dark gray), DHSs with SNVs (light gray), DHSs passing mappability filters (orange), DHSs with sufficient coverage to test for imbalance across all data sets (green) and in individual cell types or strains (blue). Counts include only autosomal DHSs. **i**. Pearson correlation of allelic ratios at adjacent SNVs broken down by distance to next SNV. Dashed line represents the median width of DHS hotspots overlapping SNVs in this study.

We identified an average of 196,276 DHS hotspots (FDR 5%) in each condition using the program hotspot2^1^, and generated master lists of DHSs for each strain/cell type combination (**Supplementary Table 2**). Hierarchical clustering showed that samples clustered by cell or tissue type, rather than by strain (**Fig. 1d**), suggesting that additional strains provide access to novel genetic diversity while demonstrating consistent cell-type specific regulatory landscapes.

To identify sites of allelic imbalance indicative of genetic differences affecting DNA accessibility, we developed a custom pipeline to filter and count reads mapping to each allele at known point variants in DHSs (Methods). The majority of SNVs were testable in only a single strain or cell/tissue type, suggesting that additional profiling is likely to yield further insights (**Fig. 1e-f**). We used a beta binomial test to determine statistically significant imbalance. We applied multiple testing correction and set a significance threshold of 10% false discovery rate (FDR) and additionally required a strong magnitude of imbalance (>70% of reads mapping to one allele). Plotting the distribution of allelic ratios confirmed that our mapping strategy was not biased towards the reference allele (**Supplementary Fig. 3**). By pooling reads from multiple samples, we assessed imbalance on aggregate, per-cell type, per-strain, and per-sample bases (**Fig. 1g**). We identified a total of 13,835 strongly imbalanced SNVs out of 357,303 SNVs tested when aggregating across all samples. The high density of variation meant that nearly all DHSs in a given cell or tissue type harbored at least one SNV, and we were able to test for imbalance at an SNV in an median of 27% DHSs per cell or tissue type (**Fig. 1h**). The more highly diverged strains contributed substantially more variants tested with only a modest reduction in mappability rate (**Fig. 1h**). Full coverage of DHSs was limited primarily by sequencing depth, suggesting that additional sequencing would yield additional power. Imbalance was less frequent at highly accessible DHSs (**Supplementary Fig. 4-5**), consistent with our previous observations of buffering of point variants at strong sites^4,5^.

In the F1 offspring of an inbred cross, each variant on a given chromosome is in perfect linkage. Thus we considered the power of our approach to detect focal alteration of individual DHSs rather than coordinately altered chromatin accessibility. By examining the co-occurrence of imbalance of nearby variants, we found that allelic ratios of nearby sites were strongly correlated only at distances less than 250 bp, well below the median width of a DHS hotspot (**Fig. 1i**). This suggests that our approach offers high resolution to identify sequence variation leading to local effects on chromatin state.

### Cellular context sensitivity

We assessed the cell-type accessibility patterns in 39 diverse cell and tissue types by the ENCODE project, all mapped in reference C57BL/6 mice^24^, and excluding liver, lung, kidney, or B cells. We categorized SNVs based on whether accessibility was higher at the reference (C56BL/6J) or the non-reference allele (**Fig. 2a**). Both sets of imbalanced SNVs showed increased cell-type selectivity with respect to SNVs not affecting accessibility. But nearly half of the non-reference higher sites had evidence for a DHS in another cell or tissue type in C56BL/6J mice, a 3-fold enrichment compared to a background set of mappable SNVs in inaccessible DNA and thus not tested for imbalance (Methods). This suggests that point changes affecting accessibility at sites with preexisting activity act more frequently by broadening DNA accessibility to other cell and tissue types, rather than de novo evolution of novel regulatory DNA. Only a minority of non-reference higher sites overlapped a DHS in a cognate cell or tissue type, suggesting the majority were qualitative creation rather than quantitatively increased accessibility (**Fig. 2b**). This cell-type specific expansion of accessibility drew broadly from other cell lineages with only moderate preference for related cell types (**Fig. 2b**).

**Fig. 2.**
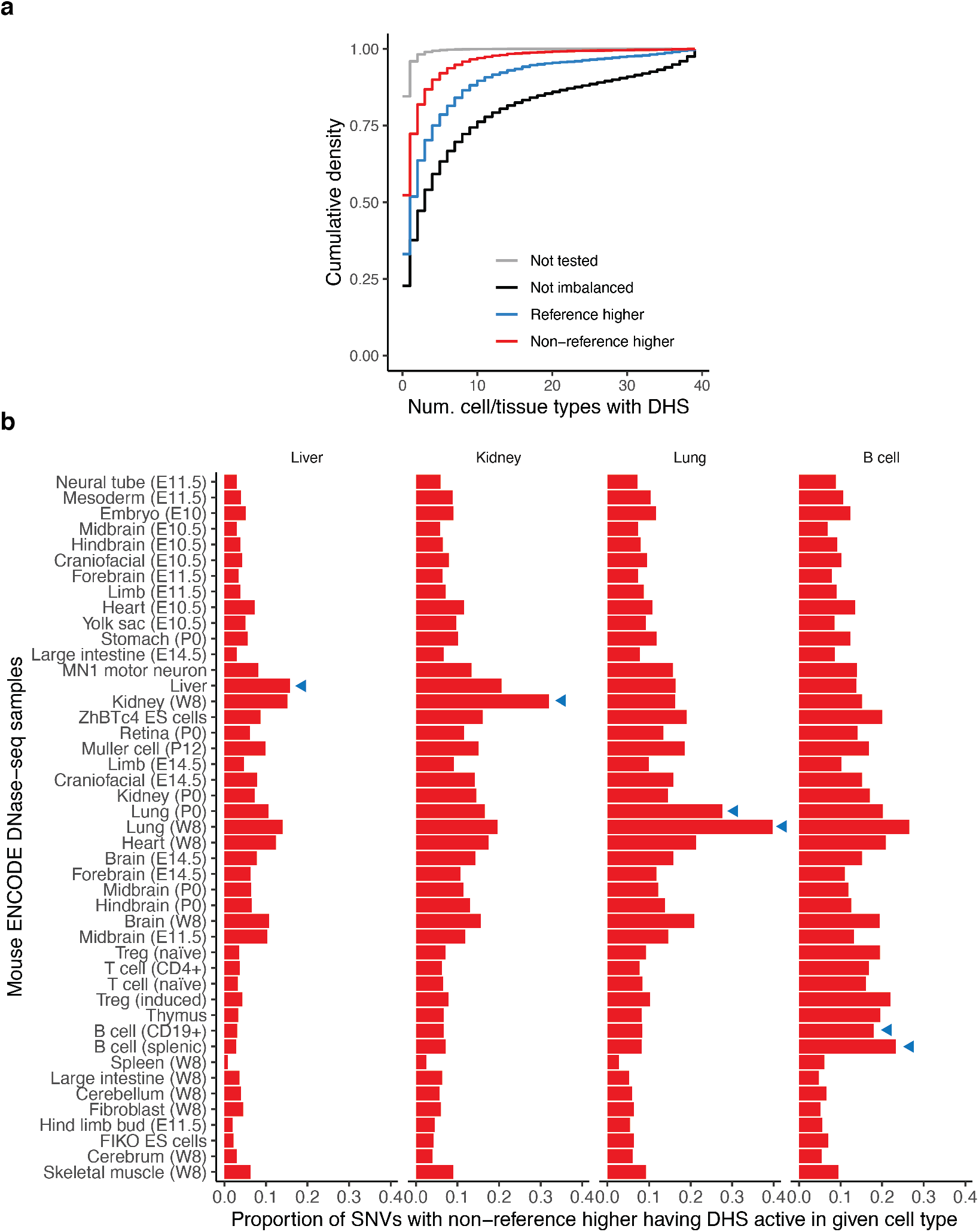
Predetermination of sites of strain-specific DNA accessibility. **a**. Cumulative density distribution of cell-type activity of DHSs measured across 39 mouse ENCODE DNase-seq samples in reference C57BL/6 mice^24^. DHSs are stratified based on whether imbalance favored the reference or non-reference allele. Not tested refers to the set of mappable SNVs not in DHSs for Liver, Kidney, Lung or B cells and therefore not tested for imbalance. **b**. Proportion of imbalanced SNVs from a given cell/tissue type that overlap DHSs from mouse ENCODE cell and tissue types (along y-axis). Cell and tissue types are ordered based on hierarchical clustering of all hotspots. Developmental timepoints for some samples are indicated in parenthesis (E, embryonic day; P, postpartum, W; adult week). Blue triangles indicate ENCODE samples matching tissues from hybrid mice in this study.

We then examined the cell-type selectivity of imbalance itself. We were able to test for imbalance per cell type (combining data from different strains) at an average of 196,276 SNVs per cell type (**Table 1**). We identified clear examples of strong imbalance across multiple strains specific to a particular cell type (**Fig. 3a-b**). In both examples, cell-type specific imbalance in one DHS was associated with a coordinate change in accessibility at a nearby DHS (**Fig. 3a-b**), though we note that it is not possible to infer the direction of causality. Overall, however, we identified a higher degree of sharing of imbalance between samples of the same cell type than from the same strain or unrelated samples (**Fig. 3c**). Pairwise comparison of different cell types showed an average of 63% sharing of imbalanced sites (1-π_0_), suggesting a high prevalence of genetic effects demonstrating cell-type context sensitivity (**Fig. 3d**).

**Table 1.**
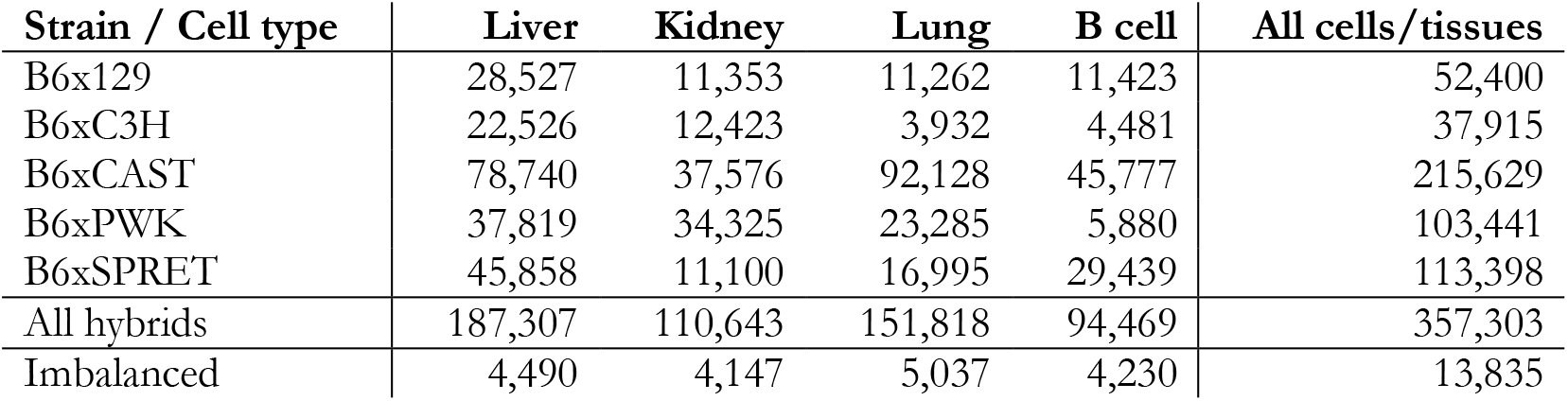
Summary of SNVs tested for imbalance per cell type/strain. Shown are counts for variants that were tested for imbalance in the per-sample, per-cell type and per-strain analyses. Imbalanced variants are shown for the per-cell type analysis in the bottom row.

| **Strain / Cell type** | **Liver** | **Kidney** | **Lung** | **B cell** | **All cells/tissues** |
| --- | --- | --- | --- | --- | --- |
| B6x129 | 28,527 | 11,353 | 11,262 | 11,423 | 52,400 |
| B6xC3H | 22,526 | 12,423 | 3,932 | 4,481 | 37,915 |
| B6xCAST | 78,740 | 37,576 | 92,128 | 45,777 | 215,629 |
| B6xPWK | 37,819 | 34,325 | 23,285 | 5,880 | 103,441 |
| B6xSPRET | 45,858 | 11,100 | 16,995 | 29,439 | 113,398 |
| All hybrids | 187,307 | 110,643 | 151,818 | 94,469 | 357,303 |
| Imbalanced | 4,490 | 4,147 | 5,037 | 4,230 | 13,835 |

**Fig. 3.**
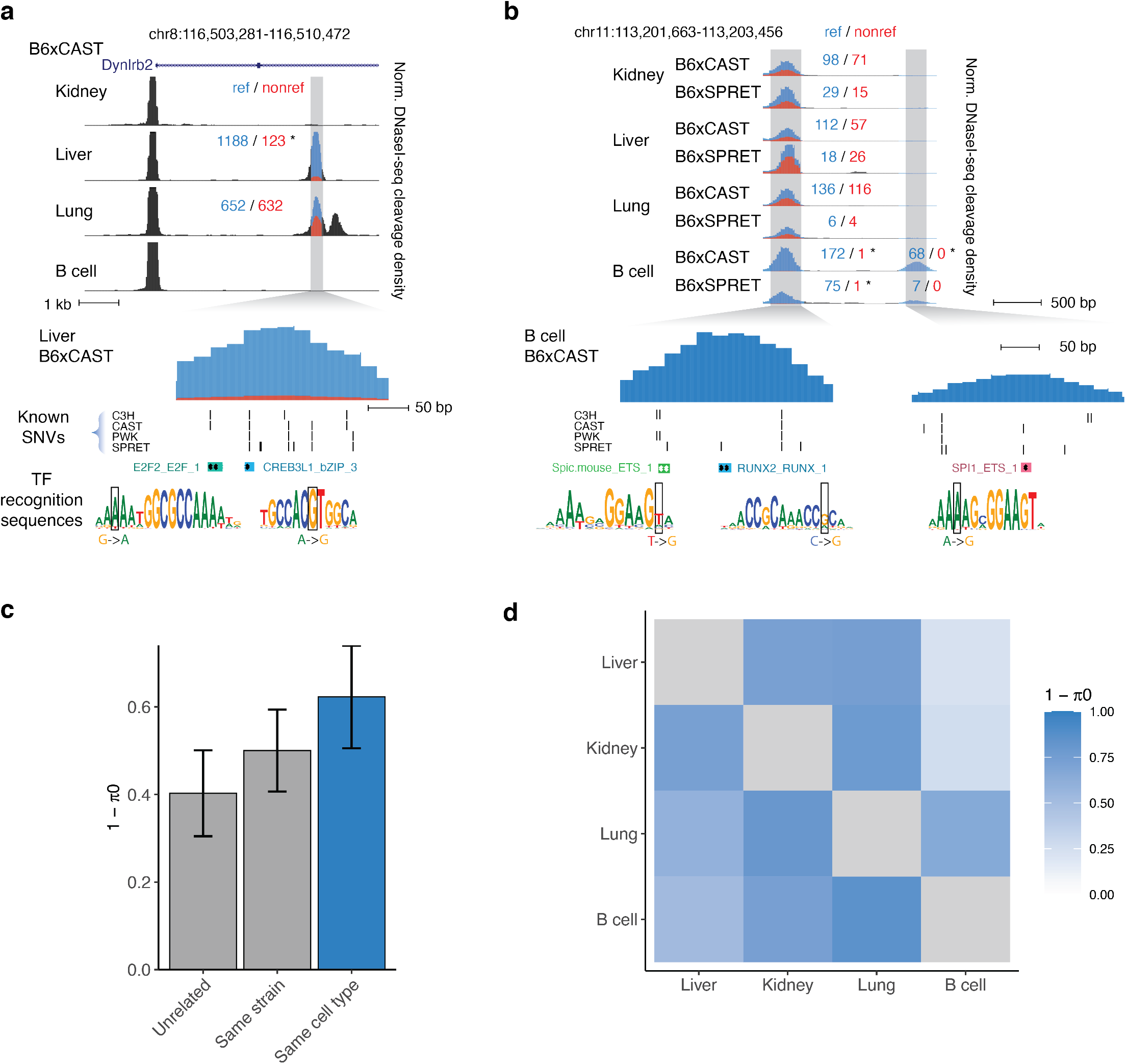
Cross-cell type analysis of allelic variation in DNA accessibility. **a-b**. Example DHSs showing cell-type specific imbalance. Normalized DNaseI cleavage density is colored by signal mapping to reference (blue) and non-reference (red) alleles based on the aggregation of informative SNVs. Counts above peaks denote sum of reads for all SNVs in region mapping to reference and non-reference alleles; * indicates statistically significant allelic imbalance. Selected TF recognition sequences overlapping imbalanced SNVs are highlighted below. **a**. Shows DHS accessible in liver and lung but with imbalance only in liver. **b**. Shows DHSs accessible in all 4 tissues (left) or specific to B cells (right); both DHSs only show imbalance in B6xCAST and B6xSPRET B cells. **c-d**. Sharing of imbalance by cell type. 1 – π_0_ represents the proportion of rejected null hypotheses by Storey’s method. **c**. Average sharing of imbalance (1 - π_0_) between samples of the same cell type vs. samples sharing only the same strain or unrelated samples (not sharing either strain or cell type). Bar height represents average of all pairwise comparisons. Error bars represent standard deviation. **d**. Pairwise sharing of imbalance between all cell or tissue types.

### TF-centric analysis of variation

We then asked to what extent variation affecting DNA accessibility in cis was linked to direct perturbation of TF recognition sequences. We scanned the mouse reference and strain-specific genomes using motif models for 2,203 TFs^4^. We found that while only a small fraction of imbalanced variation overlapped a recognition sequence for any individual TF, 61% of variation overlapped stringent motif matches (FIMO P < 10^−5^) when considering all TFs with known motifs (**Fig. 4a**). Imbalanced SNVs were found more frequently at sites of DNase I footprints, contingent on the presence of a recognizable TF recognition sequence (**Fig. 4b**). We found that aggregate imbalance was concentrated over the core positions of the motif for many key TFs (**Fig. 4c**). Sensitivity profiles for human TFs generated using previously published allelic accessibility data_4_largely resembled those generated from mouse data, although some factors such as HNF1A showed significant enrichment only in the mouse data (**Fig. 4d**).

**Fig. 4.**
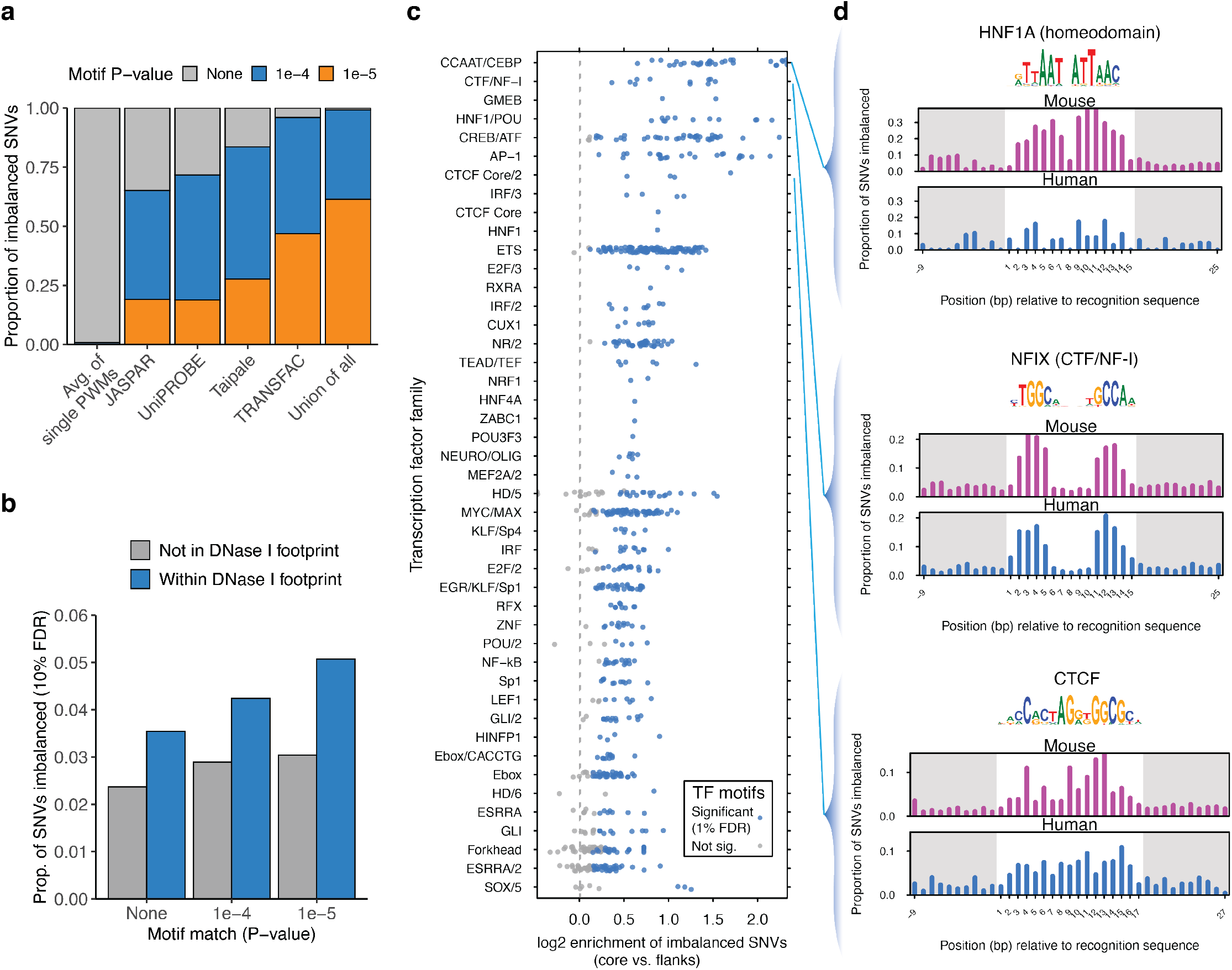
Analysis of variation affecting TF occupancy. **a**. Overlap of imbalanced SNVs with matches to TF motifs from different large-scale collections. **b**. Frequency of aggregate imbalance at SNVs overlapping TF motifs from a large-scale SELEX-seq database^48^ and DNase I footprints aggregated across all cell types. Results are stratified by FIMO value of overlapping TF motif (if any). **c**. Enrichment by TF family of imbalanced SNVs in TF core recognition sequences, relative to flanking sequence. Each point corresponds to a TF motif, grouped into TF families on the Y-axis. Shown are TF families with at least one enriched motif. **d**. TF profiles for NFIX, CTCF and HNF1A. For comparison, profiles generated from published analysis in human_4_are shown below in blue.

We next performed an analysis of cell-type specific imbalance calls at TF recognition sequences. We found higher rates of cell-type specific imbalance at sites of DNase I footprints in matching cell and tissue types, relative to unmatched cell and tissue types (**Fig. 5a**). We found that distinct TF families presented varying cell-type specific patterns of enrichment of imbalanced SNVs over their motifs (**Fig. 5b**). For example, JDP2 (AP-1) only showed enrichment in lung (**Fig. 5c**), and ETS factors showed highest enrichment in B cells (**Fig. 5d**). In both cases, no enrichment is evident when data are aggregated across multiple cell and tissue types. Other factors showed patterns of enrichment across a subset of cell types: HNF factors showed peak enrichment in liver and kidney (**Fig. 5e**), while CEBP showed enrichment in lung and liver (**Fig. 5f**). These results suggest that cell-type specific identification of imbalanced variants can yield more accurate assessment of variants affecting TF occupancy than aggregate analyses across multiple cell types.

**Fig. 5.**
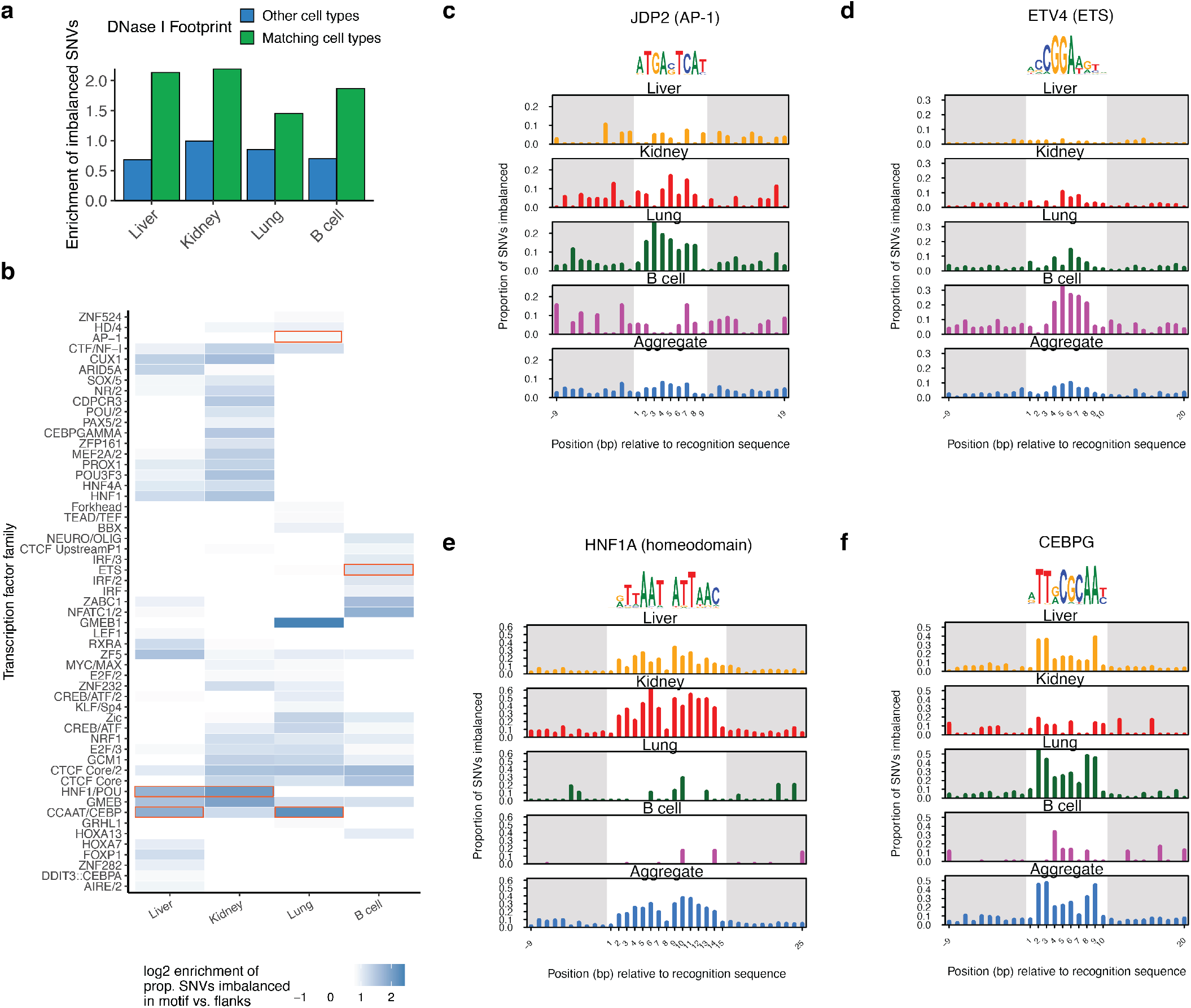
Cellular context sensitive analysis of variation affecting TF occupancy. **a**. Enrichment of imbalance called in each cell type for overlap with DNase I footprints in matching cell type (green) or in other cell types (blue). **b**. Cell-type specific enrichment of SNVs in motif for TFs. Shown are TF families with greater than twofold enrichment in at least one cell type. **c-f**. Analysis of variation affecting TF occupancy across cell/tissue types for JDP2, ETV4, HNF1A, and CEBPG motifs.

To facilitate recognition of sequence variation affecting DNA accessibility in the mouse and human genomes, we incorporated the mouse data into our Contextual Analysis of Transcription Factor Occupancy (CATO2) scoring approach^4^. CATO2 trains a logistic regression model for each TF motif on a variety of genomic annotations and TF-centric parameters. By standardizing genomic annotations between human and mouse, we directly incorporated both data sets (**Fig. 6a**). Combining the mouse and human data yielded a dramatic increase in TF families with sufficient variation (**Supplementary Table 3**). In addition to the inherent cell-type selectivity of DHS tracks, we incorporated per-cell type imbalance data in two ways (Methods): (i) TF models were trained on the subset of mouse cell types demonstrating enrichment of imbalanced SNVs over the recognition sequence (**Fig. 6b**); and (ii) a sparse generalized linear model was trained to establish cell-type specific weights for the contribution of each TF model to the overall score (**Fig. 6c**). Assessing performance on a pair of DNase-seq datasets generated in B6xCAST mouse embryonic stem cells (mESCs) (**Supplementary Table 5**) showed that CATO2 retained performance even on a completely independent validation set (**Supplementary Fig. 6)**. Furthermore, assessment of predictive performance for CTCF directly against matching ChIP-seq data showed that CATO2 scores were also predictive for allelic TF occupancy (**Supplementary Fig. 6)**. In addition, cell-type specific models showed increased predictive performance using precision-recall analysis (**Fig. 6d, Supplementary Fig. 7**). These results suggest that CATO2 provides a strong foundation for assessment of functional non-coding variation.

**Fig. 6.**
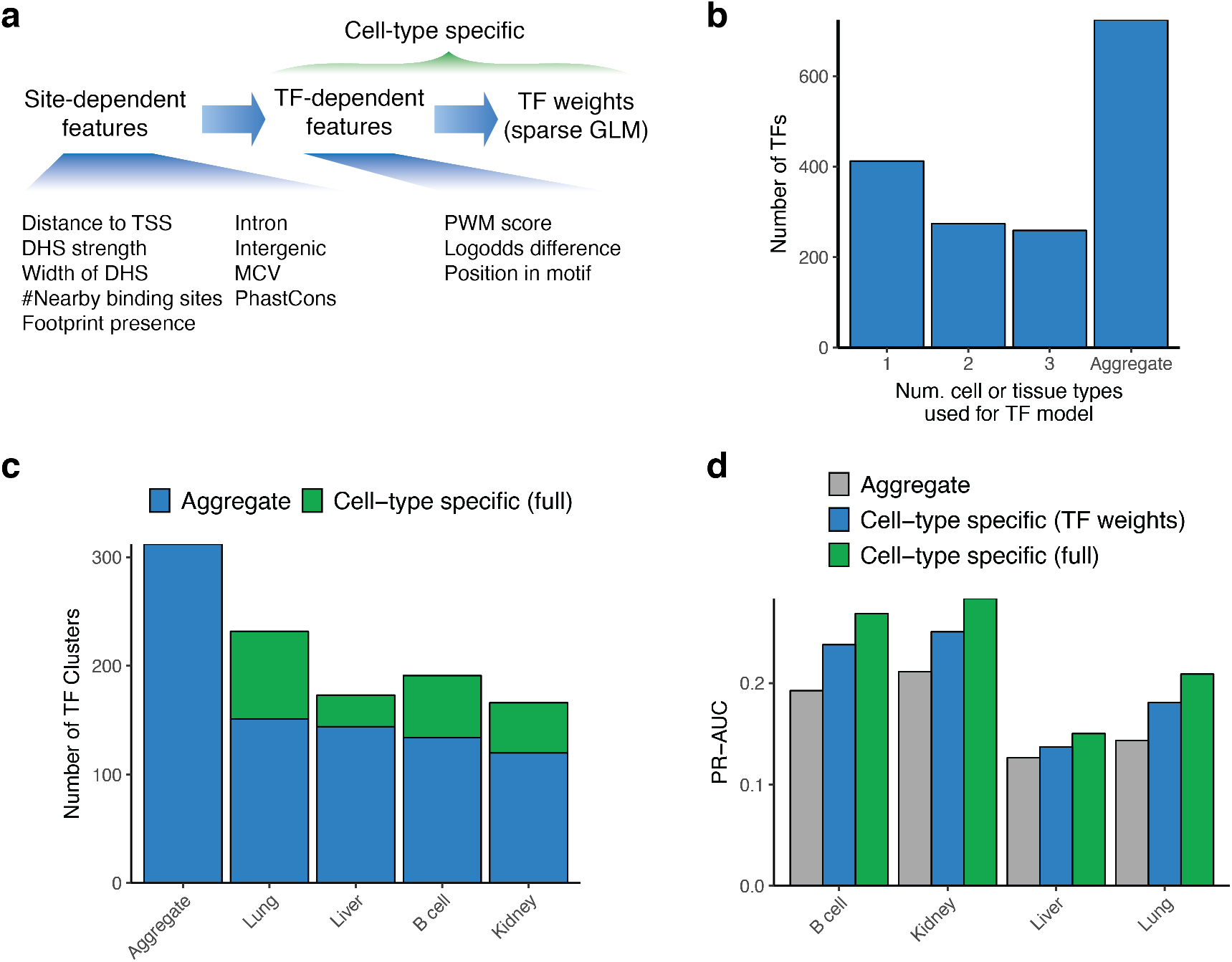
Cell-type specific prediction of variation affecting TF occupancy. a. CATO2 strategy for cell-type specific scoring of regulatory variation. b. Number of mouse cell-types used for each TF model; all TF models included human data. c. Number of unique TF clusters with non-zero coefficients in aggregate and cell-type specific CATO2 scores. TFs shared with the aggregate model are highlighted in blue. d. Area under precision-recall curves (full curves shown in **Supplementary Fig. 7**) showing performance to predict imbalanced polymorphism on SNVs tested for imbalance in individual cell types.

### Allelic effects on transcript levels

The activity of distal regulatory elements is compartmentalized and shows highly specific interactions with certain genes^25^. To examine the effect of altered accessibility on steady state transcript levels, we performed RNA-seq in a subset of matching samples (**Supplementary Table 4**). We analyzed allelic expression measured by RNA-seq using a similar pipeline to that used for the DNase-seq data (Methods). We then compared allelic accessibility at DHSs to allelic transcript levels linked to transcription start sites (TSS) within 500 kb (**Fig. 7**). We detected correlation significantly above that observed in permuted data extending distances as far as 100 kb surrounding the TSS. Maximal correlation (*R* between 0.1 and 0.2) occurred within 10 kb of the TSS, and was slightly higher downstream than upstream. Our work suggests that long-range regulatory interactions between distal accessible sites and genes are common genome-wide and are amenable to analyses using the resources and approach we have described herein.

**Fig. 7.**
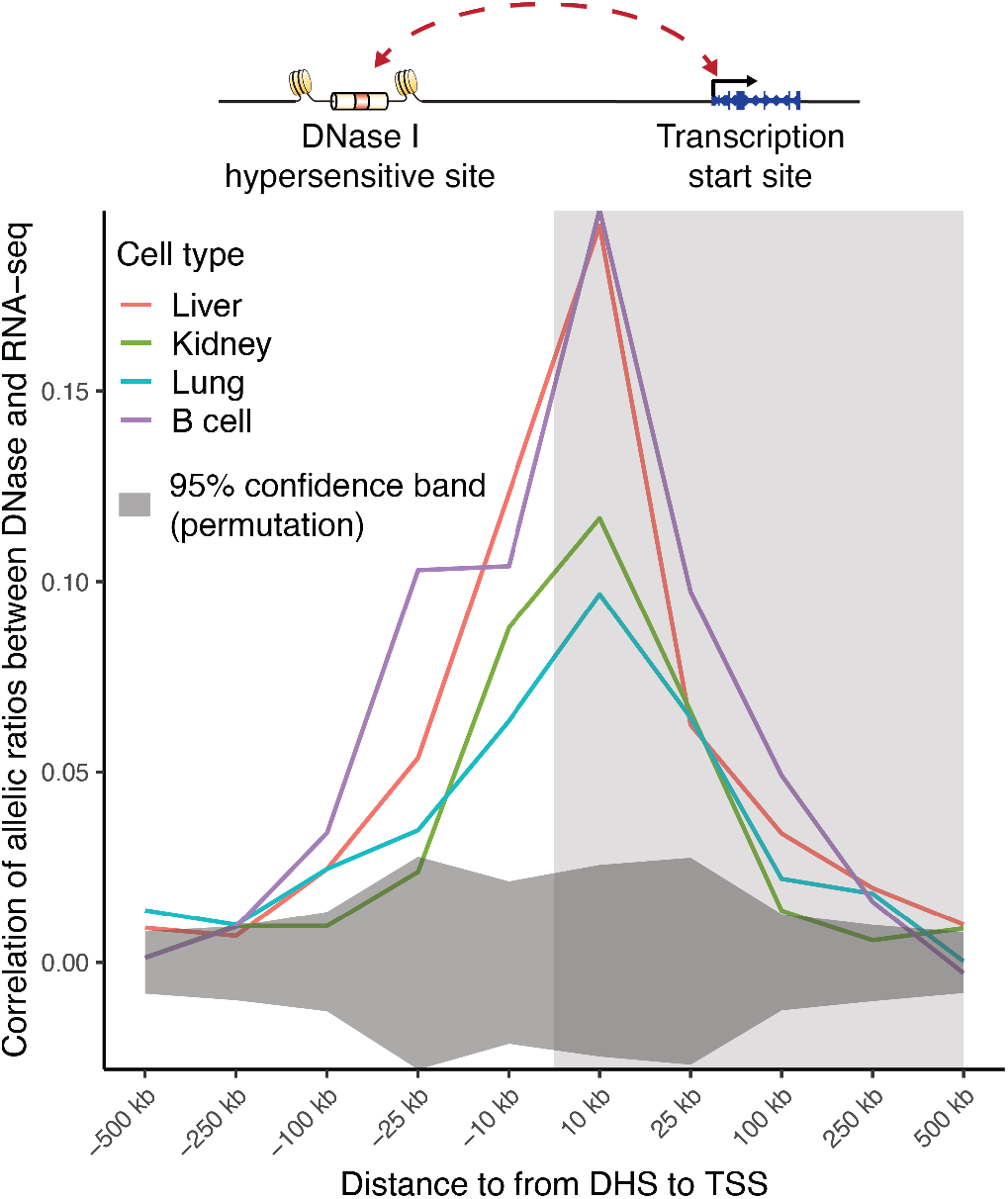
Imbalanced accessibility and transcript levels. Pearson correlation in allelic ratios between SNVs in DHSs and transcript levels broken down by distance to transcription start site (TSS). All pairs of DHSs and TSSs within 500 kb are considered. Dark gray shading at bottom indicates 95% confidence band from 1000 permutations of DHS allelic ratios among DHS-TSS pairs for each cell or tissue type. Light gray shading at right indicates that DHS lies downstream of TSS.

## Discussion

Our work shows that most cis-linked differences in DNA accessibility among diverged mouse genomes can be attributed to direct perturbation of TF recognition sites. Past reports have differed on the degree of allelic occupancy that can be linked to point changes in TF recognition sequences, ranging from 9% for NF-kB^14^ to 85% for CTCF^5^. Yet, studies of a single TF are confounded by the possibility that changes in its recognition sequence may perturb binding of other factors, either at the same site or a nearby one. By analyzing a broad set of TFs with known sequence specificities, we identify that fully 61% of imbalanced sites can be linked to changes in TF recognition sequences (**Fig. 4a**). We expect that the range of enrichment of imbalanced SNVs in TF motifs observed in **Fig. 4b** reflects both the role of cooperative binding and the accuracy of binding site recognition for individual TFs. Given the challenge of obtaining TF-specific occupancy data for all factors expressed in a given cell type, we expect that improved recognition of in vivo occupied TF binding sites from DNase I footprinting data^26,27^will be the most fruitful way to obtain further improvements in prediction performance.

Given that only a select minority of SNVs affect TF binding in a given cell type, additional large-scale analyses are needed to functionally assess noncoding variation in context. Our work shows that highly diverged mouse subspecies (including CAST/EiJ, PWK/PhJ, SPRET/EiJ) provide an efficient system for assessing regulatory variation that overcomes the low density of polymorphism in human populations. Compared to past work in human^4^, the present work required only 14% of the samples and half the sequencing depth, yet it yielded two orders of magnitude more SNVs tested for cell-type specific imbalance (avg. = 136,059 SNVs per cell type). This power enables cell-type specific analyses that uncover context-sensitive variation otherwise masked by aggregation of data across multiple cell or tissue types. The use of mice also enables ready access to a variety of cell and tissue types difficult to access in humans^21,28^. The high rate of imbalance in cell-type specific DHSs underscores the importance of robust sequencing depth across a full spectrum of cell types, and suggests that efficient generation of additional profiling data in novel cell and tissue types from these strains will efficiently increase the power of TF-centric models to recognize functional variation.

Cross-species TF-centric analysis of genomic variation overcomes the low sequence conservation of the cis-regulatory landscape^24^ by obviating the need for direct analysis of human regulatory variants at the mouse locus, and enables scalable prediction of previously unseen variation. While CATO2 presently requires cell-type specific variation data to train TF weights, inference of TF weights from other more readily available information, such as measurements of TF expression and activity, may also be possible. Such an approach could enable classification of functional regulatory variants in cell and tissue states without directly measured genetic data. Supporting this possibility, nearly half of strain-specific imbalance represented expansion of accessibility at known DHSs to a new cell or tissue type. We speculate that functional regulatory variation might most easily arise from creation of TF recognition sequences that expand the selectivity of an existing DHS by acting cooperatively with existing TF recognition sequences. It would also be straightforward to incorporate trans-regulatory differences between strains or species into future models to enable analysis of trans-regulatory effects on gene expression^29-31^.

The global correlation observed between allelic accessibility and allelic transcript levels was statistically significant but modest. Much as the majority of point variants are buffered in terms of their effect on local chromatin features^5^, enhancer networks controlling gene expression likely demonstrate a high degree of redundancy and selectivity^25,32,33^. The correlation we observe could serve as a benchmark for development of genome-wide methods to predict likely target genes of distal regulatory elements, and complements systematic locus-scale investigation of regulatory architecture using genome engineering^33,34^. Thus it is likely that further exploitation of mouse genetics will provide the substrate for more granular models of enhancer-promoter interaction.

## Methods

### Mouse husbandry

The mice used in this study were F1 hybrids of C57Bl/6J reference females with wild-derived strains 129/SvImJ (B6×129), C3H/HeJ (B6xC3H), CAST/EiJ, (B6xCAST), PWK/PhJ (B6xPWK), and SPRET/EiJ (B6xSPRET). 129/SvImJ and C3H/HeJ hybrid females were acquired from the Jackson Laboratory (8 week old, housed 4/cage). CAST/EiJ, PWK/PhJ, SPRET/EiJ inbred males were acquired from the Jackson Laboratory and were bred to C57Bl/6J female mice at FHCRC. Mice were maintained on a 12-h light, 12-h dark schedule with lights turned on at 7 a.m. The housing room was maintained at 20–24 °C with 30–70% relative humidity. Mice were housed in individually ventilated cages (Allentown) with 75 square inches of floor space and 60 air changes/hour. Biofresh cage bedding was (Absortion Corp) at 1/8 inch depth and autoclaved on site. Water and Purina 5053 (irradiated) were available *ad libitum*. Nestlet material (Envigo’s diamond twist 7979C, also irradiated) were present in each cage for enrichment. Autoclavable certified igloos (Bio-serv) were provided in some cages. Mice were housed in a barrier facility that is AAALAC accredited. Mice were sacrificed at 8 wks of age by CO_2_ asphyxiation. All work was approved by the Institutional Animal Care and Use Committee (IACUC) of the Fred Hutchinson Cancer Research Center (FHCRC).

### Nuclei isolation from mouse tissues

Solid mouse tissues were typically obtained from 4 mice sacrificed together with their tissues pooled. Whole liver (all lobes), both kidneys and all lobes of the lungs were rapidly dissected. Tissues were minced in 2 mm square pieces and resuspended in 5 mL of homogenization buffer (20 mM tricine, 25 mM D-sucrose, 15 mM NaCl, 60 mM KCl, 2 mM MgCl_2_, 0.5 mM spermidine, pH 7.8) per tissue. Nuclei were released using 5-10 strokes in a Dounce homogenizer with a loose-fitting type-A pestle and the resulting homogenate was filtered through a 120μm filter. Samples were returned to the Dounce for 5-10 strokes with a tight-fitting type-B pestle, and filtered using a 40 µm mesh filter. 5 mL of homogenate was mixed with 3 mL of 50% Optiprep solution and layered onto a 4 mL 25% - 1 mL 35% two-step Optiprep gradient and centrifuged for 20 min at 6100 x g in a swinging bucket rotor. The nuclei pellet was washed once in 10 mL of buffer A (15 mM Tris-HCl, 15 mM NaCl, 60 mM KCl, 1 mM EDTA, 0.5 mM EGTA, 0.5 mM spermidine) and resuspended at concentration of 2 × 10^6^ per mL.

Marrow was obtained from femurs of 8 week old female mice. B cells were isolated using an AutoMACS (Miltenyi Biotech) to deplete CD43 and Mac-1/CD11b markers. Cells were washed once with Dulbecco’s PBS (without MgCl_2_ or CaCl_2_). Nuclei were extracted by resuspending cells in buffer A supplemented with 0.015% detergent (IGEPAL-CA630) (Sigma) and incubating for 5-10 minutes on ice. Following incubation, the nuclei were collected by centrifugation (600 x g) and resuspended in buffer A at a concentration of 2 × 10^6^nuclei per mL.

### DNase I digestion of mouse nuclei

Fresh nuclei were incubated for 3 minutes at 37°C with limiting concentrations of the DNA endonuclease deoxyribonuclease I (DNase I) (Sigma) in buffer A supplemented with Ca^2+^. The digestion was stopped with 5X stop buffer (125 mM Tris-HCl, 250 mM NaCl, 0.25% SDS, 250 mM EDTA, 1 mM spermidine, 0.3 spermine, pH 8.0) and the samples were treated with proteinase K and RNase A. The small ‘double-hit’ fragments (<250 bp) were recovered by sucrose ultracentrifugation, end-repaired and ligated with adapters compatible with the Illumina sequencing platform. Libraries were amplified using minimal PCR cycles based on a trial qPCR amplification (8-16 cycles). A detailed protocol describing genome-wide mapping of DNase I hypersensitivity can found in ^35^.

### mESC profiling

B6xCAST mESCs were cultured as previously described in 80% 2i medium and 20% mESC medium^34^. For ChIP-seq, mESCs were crosslinked for 10 min in 1% formaldehyde and quenched in 125 mM glycine. Chromatin was sheared by Covaris LE220 Ultrasonicator (Covaris). CTCF antibody (Cell Signaling, 2899S) was conjugated to Dynabeads (M-280, Invitrogen) for 6 h at 4 °C, followed by overnight immunoprecipitation. After reversing crosslinks, immunoprecipitated DNA was treated with Proteinase K and RNase A, and purified using the DNA Clean and Concentrate-5 Kit (Zymo Research).

### Short-read sequencing and processing

DNase-seq and ChIP-seq libraries were sequenced on an Illumina HiSeq 2500 by the High-Throughput Genomics Center (University of Washington) or a NextSeq 500 (NYU Institute for Systems Genetics) in paired-end 36 bp mode.

Short reads were first trimmed to remove low-quality sequence or adapter contamination using trimmomatic v0.33^36^ with parameters ‘`TOPHRED33 ILLUMINACLIP: TruSeq3-PE-2.fa:2:5:5:1:true MAXINFO:27:0.95 TRAILING:20 MINLEN:27`’.

To reduce potential reference mapping bias, custom strain-specific genomes were created using vcf2diploid v0.26a^37^ to incorporate known^22^ point variants and insertions or deletions (REL-1505-SNPs_Indels / version 5). Chain files were created for use with the UCSC liftOver tool to enable genomic coordinate conversion between the reference and strain-specific genomes. Genomes included unscaffolded contigs and alternate sequences but not the Y chromosome.

Reads were mapped using Burrows-Wheeler Aligner (BWA) v0.7.13 to both the mouse reference assembly (GRCm38 / mm10) and the appropriate strain-specific genome with the command ‘`bwa aln -n 0.04 -l 32 -t 2 -Y`’^38^. Alignments were post-processed with a custom Python script using pysam (https://github.com/pysam-developers/pysam) to retain only properly-paired or single-end reads mapping uniquely to the autosomes and chrX with a mapping quality of at least 20. Paired end reads were required to have an inferred template length of less than 500 bp. Duplicate reads were flagged on a per-library basis using Samblaster v0.1.22 ^39^. Mapped tags were converted to BED format using awk and bedops v2.4.35^40^. DNase I hypersensitive sites were identified using hotspot2 v2.1.1^41^. Reference mm10 coordinates were used for all analyses except for read counting (which additionally relied on the strain-specific mappings).

### Assessment of allelic imbalance

Reads overlapping all known point variants were assessed for allelic imbalance at all SNVs overlapping a DNase hotspot (5% FDR) called on the aggregate of all DNase data for a given strain and cell or tissue type. Reads were extracted from DNase-seq alignments using a custom script countReads.py written using Python and pysam. The liftOver tool was used with the chain file generated by vcf2diploid to map variant coordinates from mm10 to each strain-specific genome. Reads were required to map uniquely to both mm10 and the strain-specific reference with the same mapping quality and template length. We excluded 3 bp at the 5’ end of the read to exclude any possibility of sequence-specific DNase I cut rate^42^. Only reads with a base quality >20 at the variant position were counted. Read pairs overlapping a variant were counted once. 2 additional mismatches were permitted besides the known variant. Duplicate reads passing all filters with the same 5’ position on the reference were excluded (independent of the SAM duplicate flag). Variants lying within 72 bp of a known insertion or deletion or with ≤60% of total overlapping reads passing filters were excluded from further analysis.

To minimize possible mapping bias, we generated a mappability track by mapping simulated 36-bp paired-end reads with up to 125 bp-fragment length overlapping known SNPs and including no sequencing errors. Simulated reads were mapped back to both the reference and strain-specific genomes and filtered using the approach described above. SNVs having ≤95% of simulated reads mappable were filtered out.

A background set of SNVs not tested for imbalance was identified as all mappable SNVs not overlapping a DHS in the master list or any individual condition.

Allele counts from all samples were aggregated into a single matrix and analyzed separately for per-sample, per-strain, and per-cell type imbalance. Only SNVs with at least 30 reads in one condition were retained. To account for variable sequencing depth and enrichment, we fit a beta binomial distribution for each condition using sites with >100 reads and computed *P* values against an expected 50% of reads mapping to each allele. We accounted for multiple testing using a false discovery rate (FDR) cutoff of 10% using the R package qvalue^43^. Aggregate imbalance analyses used sums of per-cell type counts.

### Transcription factor motif analysis

We scanned the reference and all strain-specific genomes using FIMO v4.10.2^44^ with TF motifs and TF clusters as in ^4^. Strain-specific motif matches were converted to mm10 coordinates using liftOver, and a non-redundant list of motif matches per-strain was created from the union of both sets.

We analyzed the intersection of SNVs tested for imbalance with these motifs. We considered motifs with a median of ≥40 SNPs per position in the motif and ≥3 positions with ≥7 significant SNPs; positions with <7 SNPs were considered missing data. For SNPs overlapping multiple matches to the same motif, we chose the best motif instance per SNP on the basis of FIMO *P* value.

### Genomic annotation

SNVs were annotated accordingly:

- Cell-type activity spectrum MCV (multi-cell verified) was computed from a set of 45 representative samples from Mouse ENCODE selected through hierarchical clustering analysis. A master list^45^was generated from these samples and MCV was scaled to 0-1 by dividing by 45.
- Footprints on mouse and human samples were called using FTD^27^.
- RefSeq Genes and CpG Islands were downloaded from the UCSC Genome Browser.

Human SNPs were annotated as in ^4^. Quantitative mouse annotations were scaled by the ratio of the mean annotation value at SNPs in mouse vs. human. Parameters were standardized to have a mean of 0 and standard deviation of 1.

### CATO2 scores

We generated CATO2 models on the combined human and mouse data as in _4_with several modifications. First, we trained a logistic model for the genomic annotations at each SNV using the glm() function in R v3.5.2:

~~~
significant ∼ MCV^2 + intron + intergenic + log(Dist. to TSS)^2 +
DHS strength^2 + log(Width of DHS) + Footprint presence + #nearby
binding sites^2 + PhastCons
~~~

Then, we trained a second glm() logistic model for each TF, which incorporated the global per-SNV score as a parameter. Imbalance was analyzed per-cell type for the mouse data and cell types demonstrating log enrichment >1 of imbalanced SNVs over the recognition sequence.

~~~
significant ∼ global.fit + log(score)^2 + logodds difference + x_2_ +
… + x_n_
~~~

Finally, we combined scores from individual TF models at each SNV using the GLMnet^46^ package to train a sparse GLM using the lasso penalty and 50-fold cross-validation with performance measured by AUC. To score human point variants, annotation values were computed and standardized as before and CATO2 scores were computed using the R function predict(type=“response”).

### Generation and analysis of RNA-seq data

Total RNA was isolated using the mirVana miRNA Isolation Kit with phenol (AM1560). Spike-in controls were mixed in (Ambion-ERCC Mix, Cat no. 4456740) and Illumina sequencing libraries were made using the RNA TruSeq Stranded total RNA (Illumina). Libraries were sequenced on an Illumina HiSeq 2500 or NextSeq by the High-Throughput Genomics Center (University of Washington) in paired-end 36 bp or 76 bp modes. Previously published data for kidney, liver, and lung B6xCAST^19^ were downloaded from the NCBI SRA repository (**Supplementary Table 4**).

Reads were mapped to the mm10 reference and strain-specific genomes in parallel using STAR v2.5.2a^47^. Counts from all non-exonic SNVs overlapping a given Gencode M10 basic level 1 and 2 protein-coding transcript were aggregated. SNVs were analyzed using same allele counting pipeline as for DNase-seq data. We assessed allelic imbalance using a beta binomial model fit at SNVs with >100 reads. We accounted for multiple testing using a false discovery rate (FDR) cutoff of 10% using the R package qvalue^43^ and additionally required >60% of reads to map to one allele. Counts were aggregated for all samples per cell type and per-DHS hotspot. A minimum of 50 total reads per transcript were required. RNA-seq imbalance data were then overlapped with per-sample DHS imbalance data.

## Data availability

Sequencing data have been deposited in the NCBI GEO repository under accession GSE156692.

## Code availability

The processing pipelines for DNase-seq and RNA-seq data are available at https://github.com/mauranolab/hybridmouse. All code for analyses herein is available upon request.

## Acknowledgements

This work was partially funded by U54HG007010 and 1S10OD017999 to J.A.S. and R35GM119703 (M.T.M). We thank Eric Rynes (Altius Institute) and Nick Vulpescu (NYU Institute for Systems Genetics) for technical assistance.

## Author Contributions

R.B., J.M.H., M.S.H., R.O., and M.T.M. performed experiments. M.A.B. and M.G. supervised mouse work. M.T.M. analyzed data. M.T.M. and J.A.S. wrote the manuscript.

## Supplemental material

## SUPPLEMENTAL FIGURES

**Supplementary Fig. 1.**
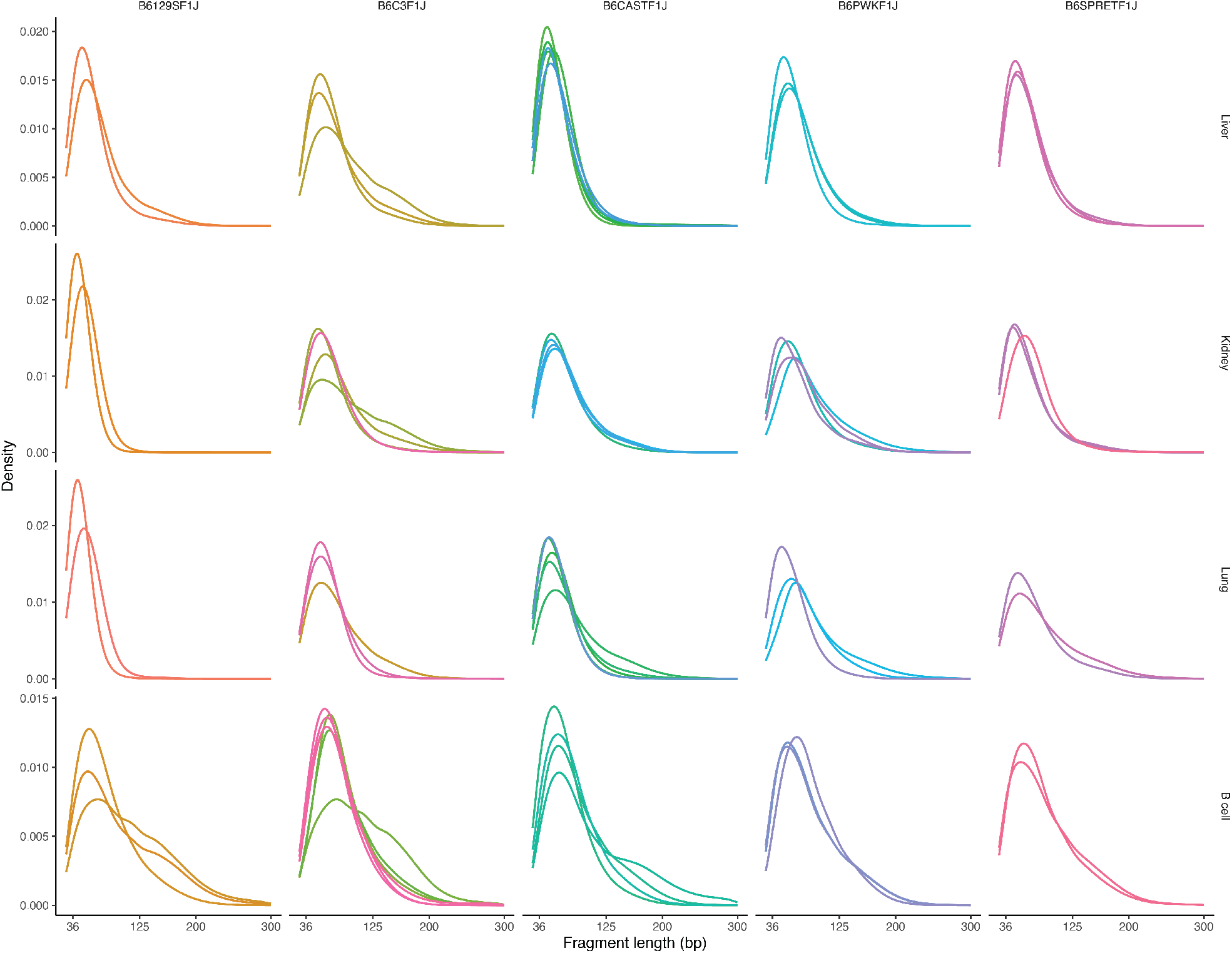
Fragment length distributions of DNase-seq data. Shown are samples passing all QC filters.

**Supplementary Fig. 2.**
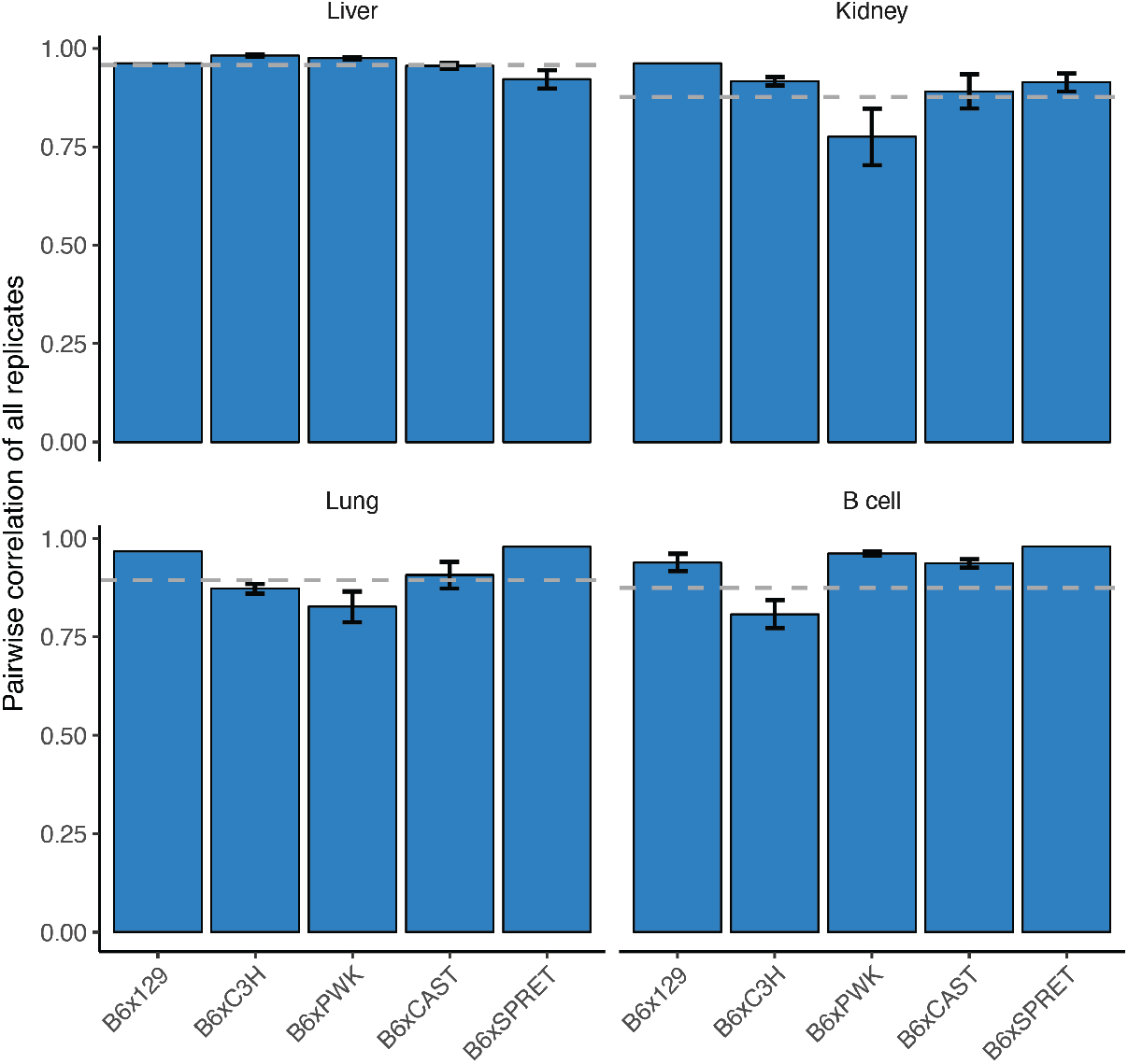
DNase-seq data replicate concordance. Average pairwise replicate concordance for each cell/tissue type and strain. Y-axis measures the average pair-wise Pearson correlation between replicates of DNase cleavage density in hotspots. Error bars represent standard deviation.

**Supplementary Fig. 3.**
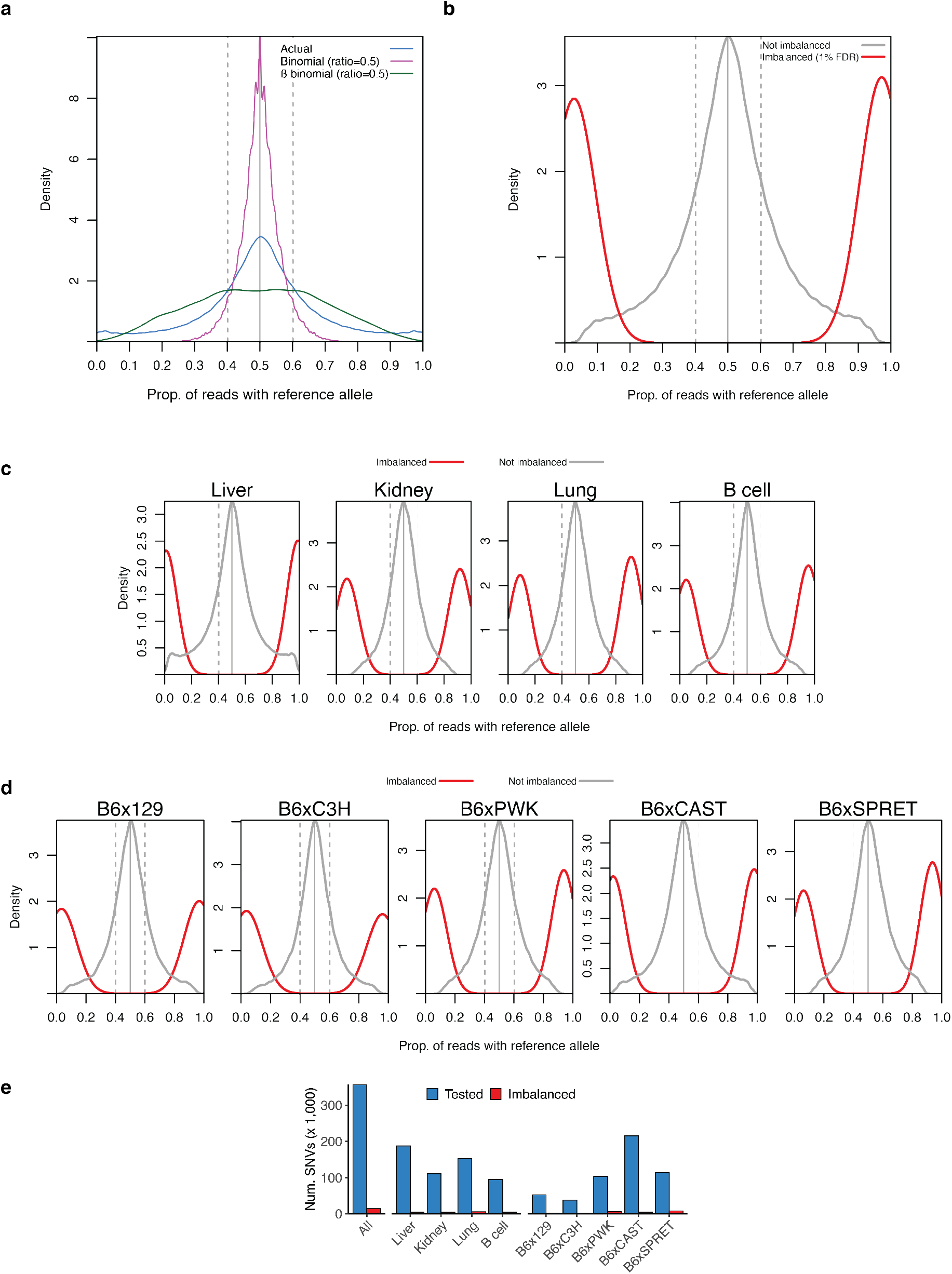
Summary of allelic imbalance analysis. **a**. Distribution of allelic ratio for actual data compared to data simulated from binomial and beta-binomial distributions. **b**-**d**. Distribution of allelic ratios for aggregate (**b**), per-cell type (**c**), per-strain (**d**) analyses. **e**. Counts of SNVs tested for imbalance (blue) and significantly imbalanced SNVs (red, FDR 10%). Counts are reported in aggregate across all data sets (left), by cell type (middle), and by parental strain (right).

**Supplementary Fig. 4.**
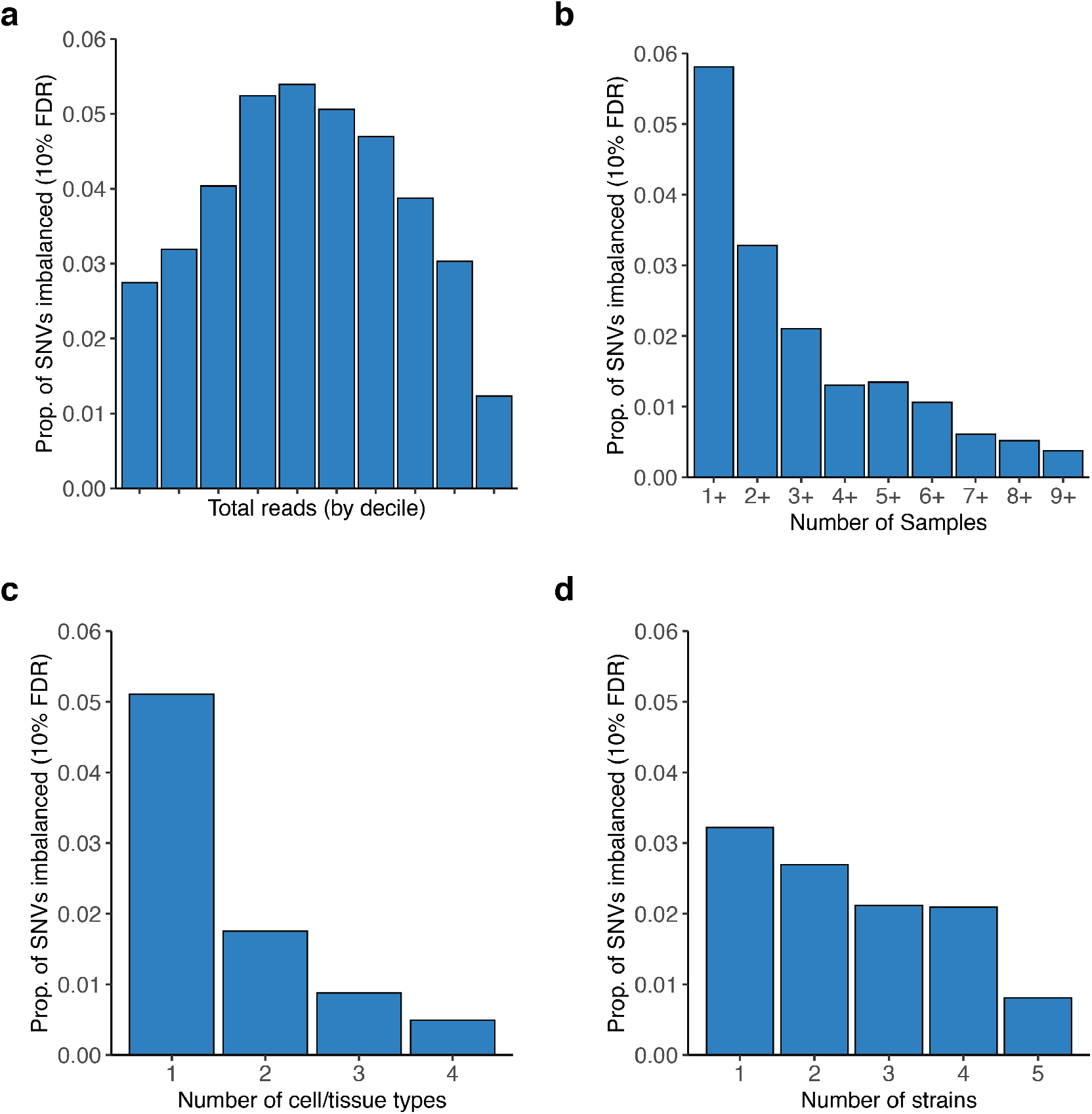
Imbalance versus read depth and number of samples. **a**. Frequency of imbalance by total reads for that SNV across all samples. **b-d**. Frequency of imbalance by the number of samples (**b**), cell/tissue types (**c**), or strains (d) in which a SNV was measured.

**Supplementary Fig. 5.**
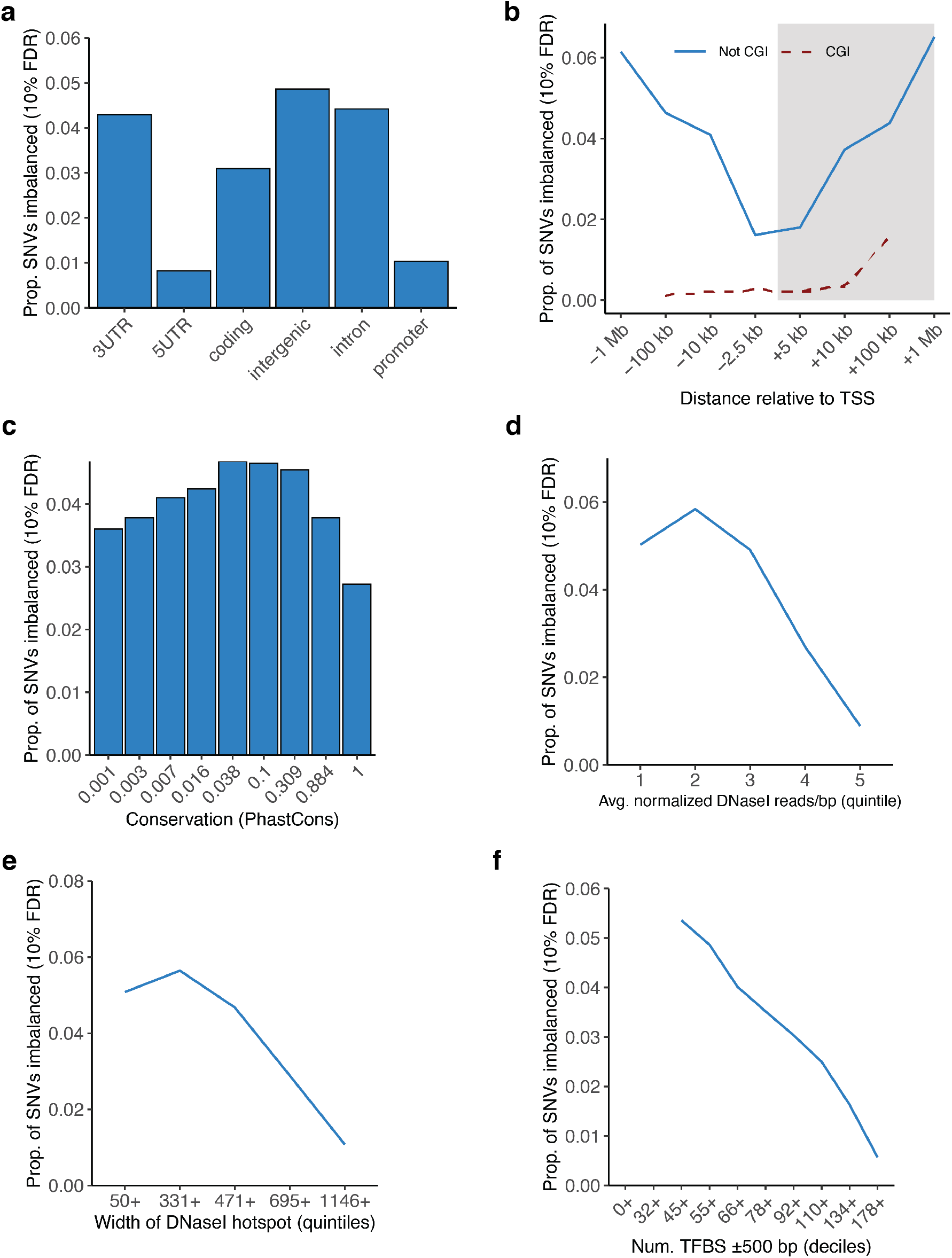
Rates of imbalance for various genomic features. Frequency of imbalance relative to **a**. genic sequence, **b**. distance to transcription start site (TSS), **c**. phylogenetic conservation (PhastCons), **d**. DHS strength, **e**. DHS hotspot width, and **f**. number of nearby TFBS in footprints.

**Supplementary Fig. 6.**
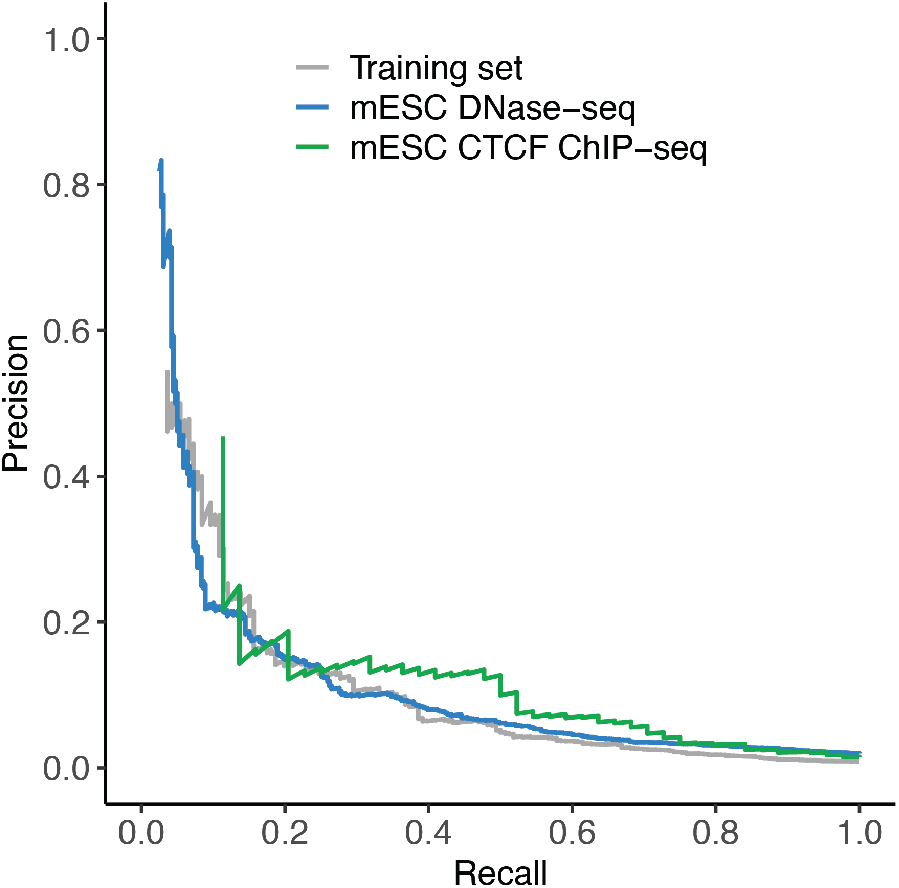
CATO2 performance on mESC validation data. Precision-recall assessment of CATO2 predictions using independently generated allelic accessibility and CTCF occupancy from mESCs (n=18,211 and 2,995 SNVs tested for imbalance, respectively). Training set performance is shown at DHSs in common with mESC data. For ChIP-seq, only predictions overlapping CTCF recognition sequences (FIMO P < 10^−4^) were assessed.

**Supplementary Fig. 7.**
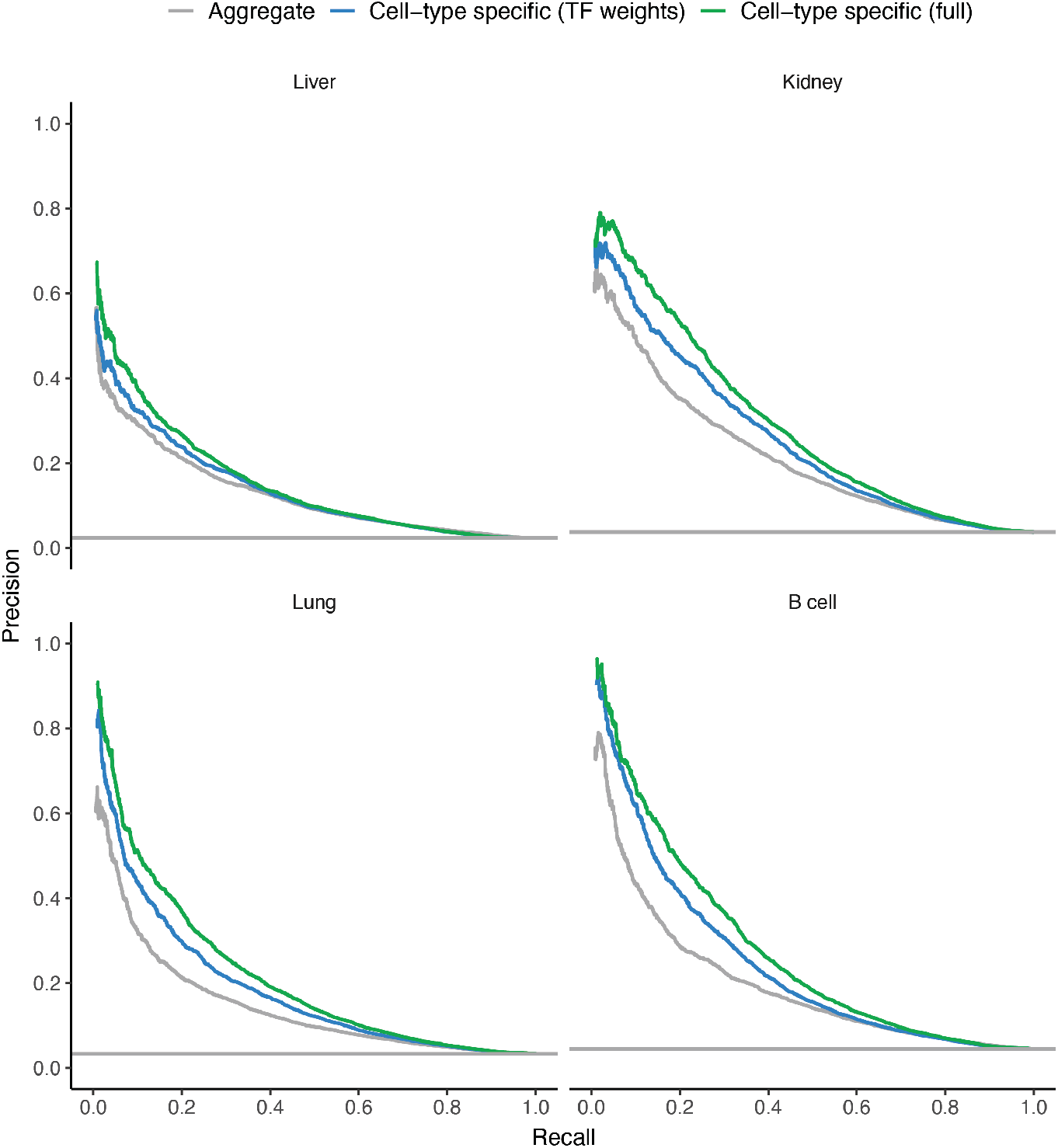
CATO2 predictive performance for variation affecting TF occupancy. Precision-recall curves showing performance to predict imbalanced polymorphism. Shown are SNVs tested for imbalance in individual cell types. Solid lines represent performance of models trained using glmnet package to have cell-type specific weights for relevant TFs. Gray lines represent performance of a random classifier based on the proportion of true positives in dataset.

## Supplementary Tables

**Supplementary Table 1.** Summary of DNase I samples in this study. Analyzed reads were required to pass all mapping filters. Read counts are in millions. Signal Portion of Tags (SPOT) scores are a measure of enrichment and refer to the proportion of reads mapping within DHS; SPOT scores are reported from hotspot V1.

| Strain | Cell/tissue type | Sample ID | # Sequenced reads (M) | # Analyzed reads (M) | # Duplicate reads (M) | SPOT |
| --- | --- | --- | --- | --- | --- | --- |
| B6x129 | B cell | DS33334 | 175 | 99 | 9 | 0.51 |
| B6x129 | B cell | DS33340 | 395 | 215 | 32 | 0.40 |
| B6x129 | B cell | DS33342 | 50 | 32 | 2 | 0.38 |
| B6x129 | Kidney | DS32758 | 176 | 121 | 49 | 0.76 |
| B6x129 | Kidney | DS32759 | 104 | 50 | 6 | 0.77 |
| B6x129 | Liver | DS32752 | 444 | 239 | 49 | 0.70 |
| B6x129 | Liver | DS32753 | 656 | 378 | 79 | 0.67 |
| B6x129 | Lung | DS32746 | 165 | 93 | 13 | 0.61 |
| B6x129 | Lung | DS32747 | 404 | 205 | 127 | 0.66 |
| B6xC3H | B cell | DS34563 | 10 | 2 | 3 | 0.81 |
| B6xC3H | B cell | DS34565 | 70 | 31 | 3 | 0.48 |
| B6xC3H | B cell | DS34569 | 9 | 2 | 3 | 0.76 |
| B6xC3H | B cell | DS38898 | 105 | 44 | 18 | 0.31 |
| B6xC3H | B cell | DS38899 | 126 | 53 | 19 | 0.36 |
| B6xC3H | B cell | DS38900 | 151 | 58 | 25 | 0.45 |
| B6xC3H | Kidney | DS33566 | 211 | 105 | 17 | 0.63 |
| B6xC3H | Kidney | DS33567 | 34 | 7 | 0 | 0.81 |
| B6xC3H | Kidney | DS33568 | 154 | 95 | 6 | 0.46 |
| B6xC3H | Kidney | DS38819 | 33 | 12 | 4 | 0.81 |
| B6xC3H | Liver | DS33560 | 247 | 98 | 7 | 0.71 |
| B6xC3H | Liver | DS33561 | 488 | 278 | 98 | 0.70 |
| B6xC3H | Liver | DS33563 | 65 | 33 | 1 | 0.59 |
| B6xC3H | Lung | DS33554 | 178 | 68 | 10 | 0.36 |
| B6xC3H | Lung | DS38811 | 131 | 63 | 49 | 0.69 |
| B6xC3H | Lung | DS38812 | 152 | 66 | 44 | 0.50 |
| B6xCAST | B cell | DS35978 | 459 | 194 | 93 | 0.79 |
| B6xCAST | B cell | DS35986 | 41 | 22 | 4 | 0.58 |
| B6xCAST | B cell | DS35992 | 63 | 27 | 6 | 0.74 |
| B6xCAST | B cell | DS35993 | 303 | 139 | 66 | 0.71 |
| B6xCAST | Kidney | DS35927 | 75 | 25 | 2 | 0.49 |
| B6xCAST | Kidney | DS36776 | 279 | 129 | 10 | 0.48 |
| B6xCAST | Kidney | DS36777 | 138 | 65 | 4 | 0.49 |
| B6xCAST | Kidney | DS36778 | 378 | 186 | 18 | 0.46 |
| B6xCAST | Liver | DS35877 | 278 | 154 | 28 | 0.69 |
| B6xCAST | Liver | DS35884 | 137 | 77 | 10 | 0.57 |
| B6xCAST | Liver | DS35889 | 77 | 30 | 3 | 0.86 |
| B6xCAST | Liver | DS35890 | 230 | 112 | 16 | 0.74 |
| B6xCAST | Liver | DS36784 | 15 | 5 | 0 | 0.76 |
| B6xCAST | Liver | DS36791 | 82 | 34 | 3 | 0.61 |
| B6xCAST | Lung | DS35897 | 84 | 44 | 3 | 0.48 |
| B6xCAST | Lung | DS35898 | 705 | 377 | 80 | 0.59 |
| B6xCAST | Lung | DS35909 | 259 | 93 | 15 | 0.45 |
| B6xCAST | Lung | DS35910 | 62 | 20 | 12 | 0.55 |
| B6xCAST | Lung | DS36795 | 24 | 7 | 0 | 0.44 |
| B6xPWK | B cell | DS36869 | 358 | 159 | 77 | 0.53 |
| B6xPWK | B cell | DS36870 | 24 | 11 | 2 | 0.46 |
| B6xPWK | B cell | DS36871 | 407 | 198 | 120 | 0.59 |
| B6xPWK | Kidney | DS36635 | 279 | 147 | 34 | 0.60 |
| B6xPWK | Kidney | DS36649 | 80 | 37 | 9 | 0.54 |
| B6xPWK | Kidney | DS37495 | 73 | 13 | 2 | 0.66 |
| B6xPWK | Kidney | DS37496 | 337 | 135 | 14 | 0.38 |
| B6xPWK | Liver | DS36636 | 64 | 32 | 2 | 0.56 |
| B6xPWK | Liver | DS36641 | 173 | 95 | 12 | 0.65 |
| B6xPWK | Liver | DS36648 | 507 | 218 | 54 | 0.86 |
| B6xPWK | Lung | DS36655 | 120 | 47 | 29 | 0.68 |
| B6xPWK | Lung | DS36657 | 527 | 202 | 77 | 0.45 |
| B6xPWK | Lung | DS37487 | 383 | 65 | 11 | 0.57 |
| B6xSPRET | B cell | DS39204 | 176 | 58 | 8 | 0.55 |
| B6xSPRET | B cell | DS39205 | 220 | 115 | 75 | 0.58 |
| B6xSPRET | Kidney | DS37590 | 225 | 58 | 4 | 0.65 |
| B6xSPRET | Kidney | DS37591 | 205 | 64 | 6 | 0.74 |
| B6xSPRET | Kidney | DS39287 | 12 | 6 | 0 | 0.56 |
| B6xSPRET | Liver | DS37603 | 292 | 139 | 15 | 0.61 |
| B6xSPRET | Liver | DS38318 | 66 | 29 | 10 | 0.81 |
| B6xSPRET | Liver | DS38327 | 178 | 71 | 36 | 0.83 |
| B6xSPRET | Lung | DS37582 | 246 | 66 | 5 | 0.48 |
| B6xSPRET | Lung | DS38311 | 197 | 104 | 7 | 0.38 |

**Supplementary Table 2.** Summary of DNase I data by strain and tissue type.

| **Strain** | **Cell/tissue type** | **Analyzed reads** | **# Biological replicates** | **# Hotspots (5% FDR)** |
| --- | --- | --- | --- | --- |
| B6x129 | B cell | 347,204,832 | 3 | 117,910 |
| B6x129 | Kidney | 170,705,656 | 2 | 238,917 |
| B6x129 | Liver | 617,160,439 | 2 | 295,565 |
| B6x129 | Lung | 298,565,608 | 2 | 261,732 |
| B6xC3H | B cell | 190,135,121 | 6 | 78,387 |
| B6xC3H | Kidney | 218,833,896 | 4 | 203,372 |
| B6xC3H | Liver | 409,883,230 | 3 | 215,510 |
| B6xC3H | Lung | 198,062,728 | 3 | 152,006 |
| B6xCAST | B cell | 394,467,062 | 4 | 156,078 |
| B6xCAST | Kidney | 404,999,564 | 4 | 228,944 |
| B6xCAST | Liver | 413,748,454 | 6 | 233,784 |
| B6xCAST | Lung | 543,296,869 | 5 | 281,978 |
| B6xPWK | B cell | 376,482,554 | 3 | 134,316 |
| B6xPWK | Kidney | 332,674,371 | 4 | 231,416 |
| B6xPWK | Liver | 345,057,236 | 3 | 242,731 |
| B6xPWK | Lung | 315,239,886 | 3 | 194,034 |
| B6xSPRET | B cell | 173,650,753 | 2 | 91,621 |
| B6xSPRET | Kidney | 127,799,573 | 3 | 186,061 |
| B6xSPRET | Liver | 239,422,480 | 3 | 211,762 |
| B6xSPRET | Lung | 170,021,189 | 2 | 169,389 |

**Supplementary Table 3.** Summary of TF models. Shown are TF motifs with enrichment of imbalanced SNVs. TF motifs were curated from multiple databases and annotated with gene name. Motifs with redundant sequence specificities by TOMTOM were identified and collapsed using a clustering approach^4^.

|  | Total TFs<br>in database | TFs overlapping sufficient variation |  |  |
| --- | --- | --- | --- | --- |
|  |  | Human (166<br>individuals and<br>116 cell types<sup>4</sup>) | Mouse (5<br>strains and<br>4 cell types) | Pooled human<br>and mouse |
| TF motifs | 2154 | 509 | 627 | 857 |
| TF genes | 695 | 268 | 335 | 430 |
| TF motifs (collapsed) | 270 | 82 | 105 | 131 |

**Supplementary Table 4.** Summary of RNA-seq samples in this study. Uniquely mapped reads were required to pass all mapping filters. Nonredundant reads exclude PCR duplicates. Read counts are in millions. Samples IDs beginning with SRR are from ^19^.

| Strain | Cell/tissue type | Sample ID | Num. pass filter alignments | Uniquely mapped reads | Nonredundant reads |
| --- | --- | --- | --- | --- | --- |
| B6xC3H | B cell | DS38895 | 199 | 173 | 94 |
| B6xC3H | Kidney | DS38815 | 128 | 113 | 77 |
| B6xC3H | Liver | DS38822 | 132 | 109 | 67 |
| B6xC3H | Lung | DS38808 | 112 | 96 | 69 |
| B6xCAST | B cell | DS35975 | 117 | 109 | 80 |
| B6xCAST | Kidney | SRR823460 | 75 | 60 | 46 |
| B6xCAST | Kidney | SRR823468 | 130 | 85 | 50 |
| B6xCAST | Liver | SRR823469 | 213 | 148 | 88 |
| B6xCAST | Liver | SRR823474 | 221 | 143 | 88 |
| B6xCAST | Lung | SRR823447 | 86 | 74 | 55 |
| B6xCAST | Lung | SRR823448 | 104 | 89 | 64 |
| B6xPWK | B cell | DS36866 | 80 | 76 | 56 |
| B6xPWK | B cell | DS37551 | 116 | 91 | 61 |
| B6xPWK | Kidney | DS37491 | 112 | 107 | 74 |
| B6xPWK | Liver | DS37504 | 124 | 100 | 55 |
| B6xPWK | Lung | DS37484 | 100 | 92 | 73 |
| B6xSPRET | B cell | DS39200 | 67 | 60 | 38 |
| B6xSPRET | Kidney | DS37587 | 103 | 98 | 67 |
| B6xSPRET | Kidney | DS38305 | 132 | 111 | 67 |
| B6xSPRET | Liver | DS37600 | 94 | 87 | 54 |
| B6xSPRET | Liver | DS38323 | 137 | 114 | 46 |
| B6xSPRET | Lung | DS37580 | 125 | 115 | 70 |
| B6xSPRET | Lung | DS38309 | 130 | 107 | 72 |

**Supplementary Table 5.**
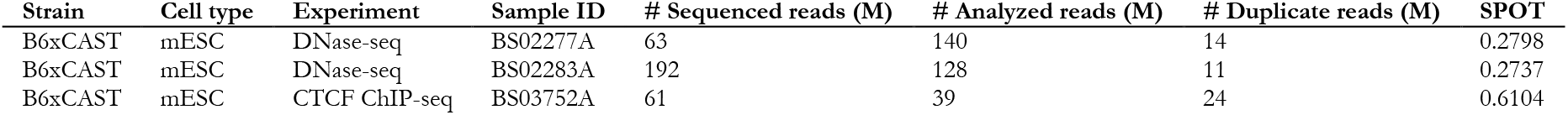
Summary of mESC samples in this study. Analyzed reads were required to pass all mapping filters. Read counts are in millions. Signal Portion of Tags (SPOT) scores are a measure of enrichment and refer to the proportion of reads mapping within DHS; SPOT scores are reported from hotspot V1.

## Supplementary Data

**Supplementary Data 1. Details of sites tested for imbalance**.

Details of 357,303 SNVs tested for imbalance, including coordinates (mm10), read counts, p-value, and aggregate and per-cell/tissue type imbalance calls.

## References

1. Thurman, R. E., Rynes, E., Humbert, R., Vierstra, J., Maurano, M. T., Haugen, E., Sheffield, N. C., Stergachis, A. B., Wang, H., Vernot, B., Garg, K., John, S., Sandstrom, R., Bates, D., Boatman, L., Canfield, T. K., Diegel, M., Dunn, D., Ebersol, A. K., Frum, T., Giste, E., Johnson, A. K., Johnson, E. M., Kutyavin, T., Lajoie, B., Lee, B.-K., Lee, K., London, D., Lotakis, D., Neph, S., Neri, F., Nguyen, E. D., Qu, H., Reynolds, A. P., Roach, V., Safi, A., Sanchez, M. E., Sanyal, A., Shafer, A., Simon, J. M., Song, L., Vong, S., Weaver, M., Yan, Y., Zhang, Z., Zhang, Z., Lenhard, B., Tewari, M., Dorschner, M. O., Hansen, R. S., Navas, P. A., Stamatoyannopoulos, G., Iyer, V. R., Lieb, J. D., Sunyaev, S. R., Akey, J. M., Sabo, P. J., Kaul, R., Furey, T. S., Dekker, J., Crawford, G. E. & Stamatoyannopoulos, J. A. The accessible chromatin landscape of the human genome. Nature 489, 75–82 (2012).

2. Maurano, M. T., Humbert, R., Rynes, E., Thurman, R. E., Haugen, E., Wang, H., Reynolds, A. P., Sandstrom, R., Qu, H., Brody, J., Shafer, A., Neri, F., Lee, K., Kutyavin, T., Stehling-Sun, S., Johnson, K., Canfield, T. K., Giste, E., Diegel, M., Bates, D., Hansen, R. S., Neph, S., Sabo, P. J., Heimfeld, S., Raubitschek, A., Ziegler, S., Cotsapas, C., Sotoodehnia, N., Glass, I., Sunyaev, S. R., Kaul, R. & Stamatoyannopoulos, J. A. Systematic localization of common disease-associated variation in regulatory DNA. Science 337, 1190–1195 (2012).

3. Palmiter, R. D. & Brinster, R. L. Germ-line transformation of mice. Annu. Rev. Genet. 20, 465–499 (1986).

4. Maurano, M. T., Haugen, E., Sandstrom, R., Vierstra, J., Shafer, A., Kaul, R. & Stamatoyannopoulos, J. A. Large-scale identification of sequence variants influencing human transcription factor occupancy in vivo. Nat. Genet. 47, 1393–1401 (2015).

5. Maurano, M. T., Wang, H., Kutyavin, T. & Stamatoyannopoulos, J. A. Widespread site-dependent buffering of human regulatory polymorphism. PLoS Genet. 8, e1002599 (2012).

6. Knight, J. C., Keating, B. J., Rockett, K. A. & Kwiatkowski, D. P. In vivo characterization of regulatory polymorphisms by allele-specific quantification of RNA polymerase loading. Nat. Genet. 33, 469– 475 (2003).

7. Degner, J. F., Pai, A. A., Pique-Regi, R., Veyrieras, J.-B., Gaffney, D. J., Pickrell, J. K., De Leon, S., Michelini, K., Lewellen, N., Crawford, G. E., Stephens, M., Gilad, Y. & Pritchard, J. K. DNase?I sensitivity QTLs are a major determinant of human expression variation. Nature 482, 390–394 (2012).

8. McDaniell, R., Lee, B.-K., Song, L., Liu, Z., Boyle, A. P., Erdos, M. R., Scott, L. J., Morken, M. A., Kucera, K. S., Battenhouse, A., Keefe, D., Collins, F. S., Willard, H. F., Lieb, J. D., Furey, T. S., Crawford, G. E., Iyer, V. R. & Birney, E. Heritable individual-specific and allele-specific chromatin signatures in humans. Science 328, 235–239 (2010).

9. Silver, L. M. Mouse Genetics: Concepts and Applications. (Oxford University Press, 1995).

10. Mouse Genome Sequencing Consortium, Waterston, R. H., Lindblad-Toh, K., Birney, E., Rogers, J., Abril, J. F., Agarwal, P., Agarwala, R., Ainscough, R., Alexandersson, M., An, P., Antonarakis, S. E., Attwood, J., Baertsch, R., Bailey, J., Barlow, K., Beck, S., Berry, E., Birren, B., Bloom, T., Bork, P., Botcherby, M., Bray, N., Brent, M. R., Brown, D. G., Brown, S. D., Bult, C., Burton, J., Butler, J., Campbell, R. D., Carninci, P., Cawley, S., Chiaromonte, F., Chinwalla, A. T., Church, D. M., Clamp, M., Clee, C., Collins, F. S., Cook, L. L., Copley, R. R., Coulson, A., Couronne, O., Cuff, J., Curwen, V., Cutts, T., Daly, M., David, R., Davies, J., Delehaunty, K. D., Deri, J., Dermitzakis, E. T., Dewey, C., Dickens, N. J., Diekhans, M., Dodge, S., Dubchak, I., Dunn, D. M., Eddy, S. R., Elnitski, L., Emes, R. D., Eswara, P., Eyras, E., Felsenfeld, A., Fewell, G. A., Flicek, P., Foley, K., Frankel, W. N., Fulton, L. A., Fulton, R. S., Furey, T. S., Gage, D., Gibbs, R. A., Glusman, G., Gnerre, S., Goldman, N., Goodstadt, L., Grafham, D., Graves, T. A., Green, E. D., Gregory, S., Guigó, R., Guyer, M., Hardison, R. C., Haussler, D., Hayashizaki, Y., Hillier, L. W., Hinrichs, A., Hlavina, W., Holzer, T., Hsu, F., Hua, A., Hubbard, T., Hunt, A., Jackson, I., Jaffe, D. B., Johnson, L. S., Jones, M., Jones, T. A., Joy, A., Kamal, M., Karlsson, E. K., Karolchik, D., Kasprzyk, A., Kawai, J., Keibler, E., Kells, C., Kent, W. J., Kirby, A., Kolbe, D. L., Korf, I., Kucherlapati, R. S., Kulbokas, E. J., Kulp, D., Landers, T., Leger, J. P., Leonard, S., Letunic, I., Levine, R., Li, J., Li, M., Lloyd, C., Lucas, S., Ma, B., Maglott, D. R., Mardis, E. R., Matthews, L., Mauceli, E., Mayer, J. H., McCarthy, M., McCombie, W. R., McLaren, S., McLay, K., McPherson, J. D., Meldrim, J., Meredith, B., Mesirov, J. P., Miller, W., Miner, T. L., Mongin, E., Montgomery, K. T., Morgan, M., Mott, R., Mullikin, J. C., Muzny, D. M., Nash, W. E., Nelson, J. O., Nhan, M. N., Nicol, R., Ning, Z., Nusbaum, C., O’Connor, M. J., Okazaki, Y., Oliver, K., Overton-Larty, E., Pachter, L., Parra, G., Pepin, K. H., Peterson, J., Pevzner, P., Plumb, R., Pohl, C. S., Poliakov, A., Ponce, T. C., Ponting, C. P., Potter, S., Quail, M., Reymond, A., Roe, B. A., Roskin, K. M., Rubin, E. M., Rust, A. G., Santos, R., Sapojnikov, V., Schultz, B., Schultz, J., Schwartz, M. S., Schwartz, S., Scott, C., Seaman, S., Searle, S., Sharpe, T., Sheridan, A., Shownkeen, R., Sims, S., Singer, J. B., Slater, G., Smit, A., Smith, D. R., Spencer, B., Stabenau, A., Stange-Thomann, N., Sugnet, C., Suyama, M., Tesler, G., Thompson, J., Torrents, D., Trevaskis, E., Tromp, J., Ucla, C., Ureta-Vidal, A., Vinson, J. P., Niederhausern Von, A. C., Wade, C. M., Wall, M., Weber, R. J., Weiss, R. B., Wendl, M. C., West, A. P., Wetterstrand, K., Wheeler, R., Whelan, S., Wierzbowski, J., Willey, D., Williams, S., Wilson, R. K., Winter, E., Worley, K. C., Wyman, D., Yang, S., Yang, S.-P., Zdobnov, E. M., Zody, M. C. & Lander, E. S. Initial sequencing and comparative analysis of the mouse genome. Nature 420, 520–562 (2002).

11. Peterson, K. R., Clegg, C. H., Huxley, C., Josephson, B. M., Haugen, H. S., Furukawa, T. & Stamatoyannopoulos, G. Transgenic mice containing a 248-kb yeast artificial chromosome carrying the human beta-globin locus display proper developmental control of human globin genes. Proceedings of the National Academy of Sciences of the United States of America 90, 7593–7597 (1993).

12. Wilson, M. D., Barbosa-Morais, N. L., Schmidt, D., Conboy, C. M., Vanes, L., Tybulewicz, V. L. J., Fisher, E. M. C., Tavaré, S. & Odom, D. T. Species-specific transcription in mice carrying human chromosome 21. Science 322, 434–438 (2008).

13. Breeze, C. E., Lazar, J., Mercer, T., Halow, J., Washington, I., Lee, K., Ibarrientos, S., Castillo, A., Neri, F., Haugen, E., Rynes, E., Reynolds, A., Bates, D., Diegel, M., Dunn, D., Kaul, R., Sandstrom, R., Meuleman, W., Bender, M. A., Groudine, M. & Stamatoyannopoulos, J. A. Atlas and developmental dynamics of mouse DNase I hypersensitive sites. bioRxiv doi:10.1093/nar/gkt1229

14. Heinz, S., Romanoski, C. E., Benner, C., Allison, K. A., Kaikkonen, M. U., Orozco, L. D. & Glass, C. K. Effect of natural genetic variation on enhancer selection and function. Nature 503, 487–492 (2013).

15. Wong, E. S., Schmitt, B. M., Kazachenka, A., Thybert, D., Redmond, A., Connor, F., Rayner, T. F., Feig, C., Ferguson-Smith, A. C., Marioni, J. C., Odom, D. T. & Flicek, P. Interplay of cis and trans mechanisms driving transcription factor binding and gene expression evolution. Nature Commun. 8, 1092 (2017).

16. Hosseini, M., Goodstadt, L., Hughes, J. R., Kowalczyk, M. S., De Gobbi, M., Otto, G. W., Copley, R. R., Mott, R., Higgs, D. R. & Flint, J. Causes and consequences of chromatin variation between inbred mice. PLoS Genet. 9, e1003570 (2013).

17. Xu, J., Carter, A. C., Gendrel, A.-V., Attia, M., Loftus, J., Greenleaf, W. J., Tibshirani, R., Heard, E. & Chang, H. Y. Landscape of monoallelic DNA accessibility in mouse embryonic stem cells and neural progenitor cells. Nat. Genet. 49, 377–386 (2017).

18. Schadt, E. E., Monks, S. A., Drake, T. A., Lusis, A. J., Che, N., Colinayo, V., Ruff, T. G., Milligan, S. B., Lamb, J. R., Cavet, G., Linsley, P. S., Mao, M., Stoughton, R. B. & Friend, S. H. Genetics of gene expression surveyed in maize, mouse and man. Nature 422, 297–302 (2003).

19. Babak, T., DeVeale, B., Tsang, E. K., Zhou, Y., Li, X., Smith, K. S., Kukurba, K. R., Zhang, R., Li, J. B., van der Kooy, D., Montgomery, S. B. & Fraser, H. B. Genetic conflict reflected in tissue-specific maps of genomic imprinting in human and mouse. Nat. Genet. 47, 544–549 (2015).

20. Crowley, J. J., Zhabotynsky, V., Sun, W., Huang, S., Pakatci, I. K., Kim, Y., Wang, J. R., Morgan, A. P., Calaway, J. D., Aylor, D. L., Yun, Z., Bell, T. A., Buus, R. J., Calaway, M. E., Didion, J. P., Gooch, T. J., Hansen, S. D., Robinson, N. N., Shaw, G. D., Spence, J. S., Quackenbush, C. R., Barrick, C. J., Nonneman, R. J., Kim, K., Xenakis, J., Xie, Y., Valdar, W., Lenarcic, A. B., Wang, W., Welsh, C. E., Fu, C.-P., Zhang, Z., Holt, J., Guo, Z., Threadgill, D. W., Tarantino, L. M., Miller, D. R., Zou, F., McMillan, L., Sullivan, P. F. & Pardo-Manuel de Villena, F. Analyses of allele-specific gene expression in highly divergent mouse crosses identifies pervasive allelic imbalance. Nat. Genet. 47, 353–360 (2015).

21. Chick, J. M., Munger, S. C., Simecek, P., Huttlin, E. L., Choi, K., Gatti, D. M., Raghupathy, N., Sven-son, K. L., Churchill, G. A. & Gygi, S. P. Defining the consequences of genetic variation on a proteome-wide scale. Nature 534, 500–505 (2016).

22. Keane, T. M., Goodstadt, L., Danecek, P., White, M. A., Wong, K., Yalcin, B., Heger, A., Agam, A., Slater, G., Goodson, M., Furlotte, N. A., Eskin, E., Nellåker, C., Whitley, H., Cleak, J., Janowitz, D., Hernandez-Pliego, P., Edwards, A., Belgard, T. G., Oliver, P. L., McIntyre, R. E., Bhomra, A., Nicod, J., Gan, X., Yuan, W., van der Weyden, L., Steward, C. A., Bala, S., Stalker, J., Mott, R., Durbin, R., Jackson, I. J., Czechanski, A. Guerra-Assunçãowi, J. A., Donahue, L. R., Reinholdt, L. G., Payseur, B. A., Ponting, C. P., Birney, E., Flint, J. & Adams, D. J. Mouse genomic variation and its effect on phenotypes and gene regulation. Nature 477, 289–294 (2011).

23. Dickinson, M. E., Flenniken, A. M., Ji, X., Teboul, L., Wong, M. D., White, J. K., Meehan, T. F., Weninger, W. J., Westerberg, H., Adissu, H., Baker, C. N., Bower, L., Brown, J. M., Caddle, L. B., Chiani, F., Clary, D., Cleak, J., Daly, M. J., Denegre, J. M., Doe, B., Dolan, M. E., Edie, S. M., Fuchs, H., Gai-lus-Durner, V., Galli, A., Gambadoro, A., Gallegos, J., Guo, S., Horner, N. R., Hsu, C.-W., Johnson, S. J., Kalaga, S., Keith, L. C., Lanoue, L., Lawson, T. N., Lek, M., Mark, M., Marschall, S., Mason, J., McElwee, M. L., Newbigging, S., Nutter, L. M. J., Peterson, K. A., Ramirez-Solis, R., Rowland, D. J., Ryder, E., Samocha, K. E., Seavitt, J. R., Selloum, M., Szoke-Kovacs, Z., Tamura, M., Trainor, A. G., Tudose, I., Wakana, S., Warren, J., Wendling, O., West, D. B., Wong, L., Yoshiki, A., International Mouse Phenotyping Consortium, Jackson Laboratory, Infrastructure Nationale PHENOMIN, Institut Clinique de la Souris (ICS), Charles River Laboratories, MRC Harwell, Toronto Centre for Phenogenomics, Wellcome Trust Sanger Institute, RIKEN BioResource Center, MacArthur, D. G., Tocchini-Valentini, G. P., Gao, X., Flicek, P., Bradley, A., Skarnes, W. C., Justice, M. J., Parkinson, H. E., Moore, M., Wells, S., Braun, R. E., Svenson, K. L., de Angelis, M. H., Hérault, Y., Mohun, T., Mallon, A.-M., Henkelman, R. M., Brown, S. D. M., Adams, D. J., Lloyd, K. C. K., McKerlie, C., Beaudet, A. L., Bucan, M. & Murray, S. A. High-throughput discovery of novel developmental phenotypes. Nature 537, 508–514 (2016).

24. Vierstra, J., Rynes, E., Sandstrom, R., Zhang, M., Canfield, T., Hansen, R. S., Stehling-Sun, S., Sabo, P. J., Byron, R., Humbert, R., Thurman, R. E., Johnson, A. K., Vong, S., Lee, K., Bates, D., Neri, F., Diegel, M., Giste, E., Haugen, E., Dunn, D., Wilken, M. S., Josefowicz, S., Samstein, R., Chang, K.-H., Eichler, E. E., De Bruijn, M., Reh, T. A., Skoultchi, A., Rudensky, A., Orkin, S. H., Papayan-nopoulou, T., Treuting, P. M., Selleri, L., Kaul, R., Groudine, M., Bender, M. A. & Stamatoyannopoulos, J. A. Mouse regulatory DNA landscapes reveal global principles of cis-regulatory evolution. Science 346, 1007–1012 (2014).

25. Spitz, F. & Furlong, E. E. M. Transcription factors: from enhancer binding to developmental control. Nat. Rev. Genet. 13, 613–626 (2012).

26. Vierstra, J. & Stamatoyannopoulos, J. A. Genomic footprinting. Nat. Methods 13, 213–221 (2016).

27. Vierstra, J., Lazar, J., Sandstrom, R., Halow, J., Lee, K., Bates, D., Diegel, M., Dunn, D., Neri, F., Haugen, E., Rynes, E., Reynolds, A., Nelson, J., Johnson, A., Frerker, M., Buckley, M., Kaul, R., Meuleman, W. & Stamatoyannopoulos, J. A. Global reference mapping of human transcription factor footprints. Nature 583, 729–736 (2020).

28. Consortium, GTEX. Genetic effects on gene expression across human tissues. Nature 550, 204–213 (2017).

29. Tian, J., Keller, M. P., Oler, A. T., Rabaglia, M. E., Schueler, K. L., Stapleton, D. S., Broman, A. T., Zhao, W., Kendziorski, C., Yandell, B. S., Hagenbuch, B., Broman, K. W. & Attie, A. D. Identification of the Bile Acid Transporter Slco1a6 as a Candidate Gene That Broadly Affects Gene Expression in Mouse Pancreatic Islets. Genetics 201, 1253–1262 (2015).

30. Brynedal, B., Choi, J., Raj, T., Bjornson, R., Stranger, B. E., Neale, B. M., Voight, B. F. & Cotsapas, C. Large-Scale trans-eQTLs Affect Hundreds of Transcripts and Mediate Patterns of Transcriptional Co-regulation. Am. J. Hum. Gen. 100, 581–591 (2017).

31. Skelly, D. A., Ronald, J. & Akey, J. M. Inherited variation in gene expression. Annu. Rev. Genomics Hum. Genet. 10, 313–332 (2009).

32. Hong, J.-W., Hendrix, D. A. & Levine, M. S. Shadow enhancers as a source of evolutionary novelty. Science 321, 1314 (2008).

33. Dickel, D. E., Ypsilanti, A. R., Pla, R., Zhu, Y., Barozzi, I., Mannion, B. J., Khin, Y. S., Fukuda-Yuzawa, Y., Plajzer-Frick, I., Pickle, C. S., Lee, E. A., Harrington, A. N., Pham, Q. T., Garvin, T. H., Kato, M., Osterwalder, M., Akiyama, J. A., Afzal, V., Rubenstein, J. L. R., Pennacchio, L. A. & Visel, Ultraconserved Enhancers Are Required for Normal Development. Cell 172, 491–499.e15 (2018).

34. Brosh, R. A versatile platform for locus-scale genome rewriting. Proceedings of the National Academy of Sciences of the United States of America 1–13 doi:10.1073/pnas.2023952118/-/DCSupplemental

35. John, S., Sabo, P. J., Canfield, T. K., Lee, K., Vong, S., Weaver, M., Wang, H., Vierstra, J., Reynolds, P., Thurman, R. E. & Stamatoyannopoulos, J. A. Genome-scale mapping of DNase I hypersensitivity. Curr Protoc Mol Biol Chapter 27, Unit 21.27 (2013).

36. Bolger, A. M., Lohse, M. & Usadel, B. Trimmomatic: a flexible trimmer for Illumina sequence data. Bioinformatics 30, 2114–2120 (2014).

37. Rozowsky, J., Abyzov, A., Wang, J., Alves, P., Raha, D., Harmanci, A., Leng, J., Bjornson, R., Kong, Y., Kitabayashi, N., Bhardwaj, N., Rubin, M., Snyder, M. & Gerstein, M. AlleleSeq: analysis of allele-specific expression and binding in a network framework. Mol. Syst. Biol. 7, 522 (2011).

38. Li, H. & Durbin, R. Fast and accurate short read alignment with Burrows-Wheeler transform. Bioinformatics 25, 1754–1760 (2009).

39. Faust, G. G. & Hall, I. M. SAMBLASTER: fast duplicate marking and structural variant read extraction. Bioinformatics 30, 2503–2505 (2014).

40. Neph, S., Kuehn, M. S., Reynolds, A. P., Haugen, E., Thurman, R. E., Johnson, A. K., Rynes, E., Maurano, M. T., Vierstra, J., Thomas, S., Sandstrom, R., Humbert, R. & Stamatoyannopoulos, J. A. BEDOPS: high-performance genomic feature operations. Bioinformatics 28, 1919–1920 (2012).

41. John, S., Sabo, P. J., Thurman, R. E., Sung, M.-H., Biddie, S. C., Johnson, T. A., Hager, G. L. & Stamatoyannopoulos, J. A. Chromatin accessibility pre-determines glucocorticoid receptor binding patterns. Nat. Genet. 43, 264–268 (2011).

42. Lazarovici, A., Zhou, T., Shafer, A., Dantas Machado, A. C., Riley, T. R., Sandstrom, R., Sabo, P. J., Lu, Y., Rohs, R., Stamatoyannopoulos, J. A. & Bussemaker, H. J. Probing DNA shape and methylation state on a genomic scale with DNase I. Proceedings of the National Academy of Sciences of the United States of America 110, 6376–6381 (2013).

43. Storey, J. D. & Tibshirani, R. Statistical significance for genomewide studies. Proceedings of the National Academy of Sciences of the United States of America 100, 9440–9445 (2003).

44. Grant, C. E., Bailey, T. L. & Noble, W. S. FIMO: scanning for occurrences of a given motif. Bioinformatics 27, 1017–1018 (2011).

45. Meuleman, W., Muratov, A., Rynes, E., Halow, J., Lee, K., Bates, D., Diegel, M., Dunn, D., Neri, F., Teodosiadis, A., Reynolds, A., Haugen, E., Nelson, J., Johnson, A., Frerker, M., Buckley, M., Sandstrom, R., Vierstra, J., Kaul, R. & Stamatoyannopoulos, J. Index and biological spectrum of human DNase I hypersensitive sites. Nature 584, 244–251 (2020).

46. Friedman, J., Hastie, T. & Tibshirani, R. Regularization Paths for Generalized Linear Models via Coordinate Descent. J Stat Softw 33, 1–22 (2010).

47. Dobin, A., Davis, C. A., Schlesinger, F., Drenkow, J., Zaleski, C., Jha, S., Batut, P., Chaisson, M. & Gingeras, T. R. STAR: ultrafast universal RNA-seq aligner. Bioinformatics 29, 15–21 (2013).

48. Jolma, A., Yan, J., Whitington, T., Toivonen, J., Nitta, K. R., Rastas, P., Morgunova, E., Enge, M., Taipale, M., Wei, G., Palin, K., Vaquerizas, J. M., Vincentelli, R., Luscombe, N. M., Hughes, T. R., Lemaire, P., Ukkonen, E., Kivioja, T. & Taipale, J. DNA-Binding Specificities of Human Transcription Factors. Cell 152, 327–339 (2013).

